# Enhancer RNAs predict enhancer-gene regulatory links and are critical for enhancer function in neuronal systems

**DOI:** 10.1101/270967

**Authors:** Nancy V. N. Carullo, Robert A. Phillips, Rhiana C. Simon, Salomon A. Roman Soto, Jenna E. Hinds, Aaron J. Salisbury, Jasmin S. Revanna, Kendra D. Bunner, Lara Ianov, Faraz A. Sultan, Katherine E. Savell, Charles A. Gersbach, Jeremy J. Day

## Abstract

Genomic enhancer elements regulate gene expression programs important for neuronal fate and function and are implicated in brain disease states. Enhancers undergo bidirectional transcription to generate non-coding enhancer RNAs (eRNAs). However, eRNA function remains controversial. Here, we combined ATAC-Seq and RNA-Seq datasets from three distinct neuronal culture systems in two activity states, enabling genome-wide enhancer identification and prediction of putative enhancer-gene pairs based on correlation of transcriptional output. Notably, stimulus-dependent enhancer transcription preceded mRNA induction, and CRISPR-based activation of eRNA synthesis increased mRNA at paired genes, functionally validating enhancer-gene predictions. Focusing on enhancers surrounding the *Fos* gene, we report that targeted eRNA manipulation bidirectionally modulates *Fos* mRNA, and that *Fos* eRNAs directly interact with the histone acetyltransferase domain of the enhancer-linked transcriptional co-activator CBP. Together, these results highlight the unique role of eRNAs in neuronal gene regulation and demonstrate that eRNAs can be used to identify putative target genes.

---

TO ORCHESTRATE the precise gene expression patterns that give rise to the phenotypic and functional diversity of complex biological systems, mammalian genomes utilize millions of regulatory elements known as enhancers. Enhancers, often located many kilobases from genes that they regulate, direct transcriptional dynamics at linked genes by activation of proximal gene promoters (Heinz et al., 2015; Li et al., 2016; Wang et al., 2011). Enhancer-promoter interactions help to ensure cell- and tissue-specific gene expression profiles in the brain, defining which genes can be turned on during neuronal specification and which genes remain accessible in adult neurons (Furlong & Levine, 2018; Gray et al., 2015; Hamilton et al., 2019; Hauberg et al., 2018; Nord et al., 2013; Robson, Ringel, & Mundlos, 2019). In addition to regulating neuronal development, enhancer regions direct activity- and experience-dependent gene expression programs required for neuronal plasticity, memory formation, and behavioral adaptation to environmental stimuli (Chen et al., 2019; Gallegos, Chan, Chen, & West, 2018; Joo, Schaukowitch, Farbiak, Kilaru, & Kim, 2016; Kim et al., 2010; Malik et al., 2014; Schaukowitch et al., 2014; Telese et al., 2015; Baizabal et al., 2018; Tyssowski et al., 2018). Moreover, the majority of DNA sequence variants that possess a causal relationship to neuropsychiatric disease and intellectual disability fall in non-coding regions of DNA (Network and Pathway Analysis Subgroup of Psychiatric Genomics, 2015; Davidson et al., 2011; Dong et al., 2018; Eckart et al., 2016; Edwards et al., 2012; Y. U. Inoue & Inoue, 2016; Schizophrenia Working Group of the Psychiatric Genomics, 2014; Sanchez-Mut et al., 2018; Vermunt et al., 2014; Voisin et al., 2015; Yao et al., 2015), and the association between these polymorphisms and altered enhancer function is becoming increasingly clear. Thus, understanding how genomic enhancers regulate individual genes in neuronal systems is critical for unraveling transcriptional contributions to brain health and disease.

Recent advances in DNA sequencing have revealed that the transcriptional landscape of all mammalian organisms is far more complex than previously appreciated. In contrast to earlier predictions, a significant fraction of mammalian genomes is transcribed into non-coding RNAs, which include long non-coding RNAs (lncRNAs; generally defined as non-coding RNAs longer than 200 nucleotides) (Djebali et al., 2012; Hangauer et al., 2013; Quinn and Chang, 2016). Much of this lncRNA landscape is composed of enhancer regions which undergo bidirectional, RNA polymerase II (RNAP2)-dependent transcription to yield enhancer RNAs (eRNAs) that are generally not spliced or polyadenylated (Arner et al., 2015; Gray et al., 2015; Kim et al., 2015; Kim et al., 2010; Kim and Shiekhattar, 2015). Critically, RNA synthesis from enhancers that regulate cellular differentiation and stimulus-dependent genes precedes mRNA synthesis from these genes (Arner et al., 2015). eRNA synthesis also precedes important chromatin remodeling events that are generally used to identify enhancers (Kaikkonen et al., 2013). Even though it is unclear whether physical enhancer-promoter interactions are required for enhancer function (Fulco et al., 2019; Ghavi-Helm, Jankowski, Meiers, Viales, & Korbel, 2019; Nott et al., 2019), eRNA transcription from enhancers is highly correlated with overall enhancer activity and the presence of enhancer-promoter loops (Li et al., 2016; Sanyal et al., 2012). In neuronal systems, eRNAs arising from activity-dependent enhancers are pervasively transcribed in response to neuronal activation, plasticity-inducing stimulation, and behavioral experience (Joo et al., 2016; Kim et al., 2010; Malik et al., 2014; Schaukowitch et al., 2014; Telese et al., 2015), providing a key link between enhancers and the downstream gene expression programs that regulate brain function.

Although recent reports suggest a functional role for eRNA regulation of enhancer states, the specific nature of this role is controversial. Here, we combined genome-wide identification of regions of open chromatin with RNA-seq to investigate eRNA transcription from multiple neuronal populations in two distinct activity states. The resulting datasets enabled us to leverage variability in eRNA transcription across samples to predict functional eRNA-mRNA pairs. This approach confirms a close relationship between population-specific and activity-dependent eRNA transcription and expression of paired genes. Using enhancer-targeted CRISPR activation (CRISPRa), we validate selected enhancer-gene pairs, and demonstrate that eRNA transcription precedes mRNA induction at these loci. Next, we examined the function of specific eRNAs from well-characterized enhancers near the *Fos* gene. This immediate early gene (IEG) is broadly responsive to neuronal activity in the brain, and enhancers at this gene contribute to distinct activity-dependent induction dynamics of *Fos* mRNA (Fleischmann et al., 2003; Joo et al., 2016; Kim et al., 2010; Malik et al., 2014; Savell et al., 2016; Zovkic et al., 2014). Intriguingly, we demonstrate eRNAs from a distal *Fos* enhancer are both necessary and sufficient for induction of *Fos* mRNA, and that eRNAs interact with CREB-binding protein (CBP), an enhancer-linked histone acetyltransferase. Together, these findings provide novel convergent evidence for a key role of eRNAs in neuronal gene regulation, and demonstrate the utility of using eRNA transcript abundance to predict global, stimulus-dependent, and region-selective enhancer-gene pairs across the genome.

## RESULTS

### Chromatin accessibility and transcription predict enhancer location, activity state, and target genes

To map enhancers genome-wide for three different brain regions (primary neuronal cultures generated from cortex, hippocampus, and striatum), we took advantage of the fact that active enhancers are associated with an open chromatin structure and are bidirectionally transcribed. We first identified 191,857 regions of open chromatin (ROCs) by generating Assay for Transposase-Accessible Chromatin sequencing (ATAC-Seq) libraries from each primary neuronal culture system. Consistent with common cutoffs used to dissociate enhancers from more proximal promoters, we filtered for intergenic ROCs (iROCs) that fell outside of canonical gene boundaries and >1kb away from annotated transcription start sites. Next, we capitalized on the characteristic bidirectionality of enhancer transcription and incorporated direction-specific total RNA-seq datasets from the same neuronal culture systems to identify bidirectionally transcribed regions of open chromatin, identifying 28,492 transcribed putative enhancers (TAPEs) (**Fig. 1A, top**; **Supplementary Data Table 1**). Confirming our pipeline, these bidirectionally transcribed regions exhibited high chromatin accessibility and dual peaks of RNA expression from each strand marking TAPE centers (**Fig. 1A, bottom**). Identified TAPEs exhibited characteristic patterns of transcription, chromatin landscape, and high correlations to potential target genes as shown in the representative examples at the *Klf4* and *Sik1* loci (**Fig. 1B**). To further validate our enhancer identification pipeline, we capitalized on publicly available ENCODE datasets from postnatal day zero (P0) mouse forebrain for the major histone modifications commonly used for enhancer identification, including H3K4me1, H3K4me3, and H3K27ac (marks for active or poised enhancers; Li et al., 2016, **Fig. 1C**). TAPE centers were enriched for the histone modifications H3K4me1, H3K4me3, and H3K27ac (but not the repressive H3K27me3 modification), as well as the enhancer-linked chromatin looping factor CTCF (CCCTC-binding factor) motifs. Additionally, TAPE regions exhibited enhanced sequence conservation (PhastConsElements20way) compared to surrounding regions.

**Figure 1.**
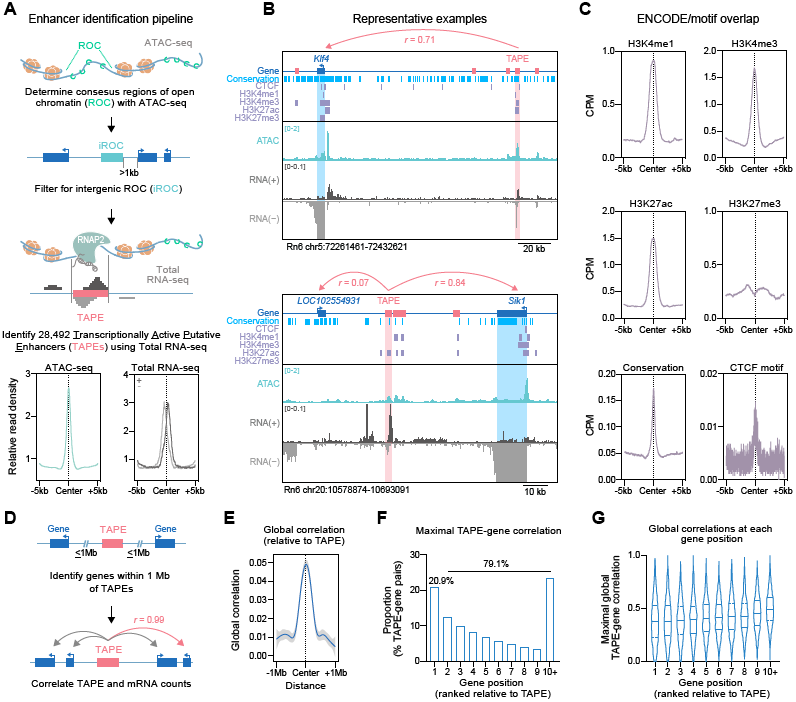
Genome-wide characterization of enhancers and eRNAs. **A**, Analysis pipeline for localization and quantification of transcriptionally active putative enhancers (TAPEs). ATAC-seq datasets were generated using cultured cortical, hippocampal, and striatal rat neurons and used to identify regions of open chromatin (ROCs). ROCs were filtered to capture intergenic regions at least 1kb from annotated genes (iROCs). Total RNA-seq data from the same culture systems was used to identify 28,492 bidirectionally transcribed intergenic ROCs and termed TAPEs. TAPEs are characterized by an enrichment of ATAC-seq and bidirectional RNA-seq reads at TAPE centers. **B**, Genome browser tracks showing ATAC-seq signal and total RNA expression at two example regions (*Klf4* and *Sik1*), relative to tracks marking conserved DNA elements (phastConsElements20way), CTCF motifs, and enhancer-linked histone modifications. **C**, TAPEs exhibit higher densities of H3K4me1, H3K4me3, H3K27ac, sequence conservation, and CTCF motifs, and decreased H3K27me3 compared to surrounding regions. Chromatin immunoprecipitation datasets were obtained from the mouse forebrain at postnatal day zero (ENCODE project) and lifted over to the rat Rn6 genome assembly. **D**, TAPE-gene pairs were determined by correlations of eRNA and mRNA levels at genes within a 1 Mb distance cutoff. **E**, On average, TAPEs and closer genes show higher correlation values. **F**, Only 20.9% of TAPEs have their maximal gene correlation at the closest gene, while the remaining 79.1% show higher correlations to genes at more distal positions. **G**, For pairs with highest global TAPE-gene correlation, correlation strength does not decrease with gene distance from TAPE.

While many pipelines have been developed to enable putative enhancer identification, the ability to predict which gene or genes are controlled by specific enhancers has remained challenging and often requires many overlapping ChIP-seq, ATAC-seq, and HiC-seq datasets for accurate prediction within a cell type. Here, we leveraged the inherent variability in eRNA levels across identified TAPEs to construct pairwise correlation matrices with mRNA estimates from all annotated protein-coding genes falling within 1 Mbp of TAPE boundaries. This strategy makes the simple assumption that enhancers and linked genes will correlate in their transcriptional output across different cell classes and/ or activity states. By filtering positively correlated, high-confidence TAPE-gene pairs from 433,416 TAPE-gene matches, we defined predicted pairs for further investigation and functional validation. Globally, identified TAPEs correlated more strongly with proximal genes than with distal genes, although average correlations were relatively weak (r < 0.05; **Fig. 1E, F**). Likewise, in contrast to traditional enhancer-gene pair prediction pipelines that simply annotate the nearest gene to an identified enhancer, our pipeline demonstrated that only 20.9% of TAPEs exhibited the strongest correlation with the closest gene, whereas 79.1% of all maximal enhancer-gene correlations occurred between TAPEs and more distal genes (ranked gene position; **Fig. 1F**). Importantly, the maximal correlation value of top TAPE-gene pairs did not decrease at more distal target genes relative to their respective TAPE (**Fig. 1G**), suggesting the presence of long-range enhancer-gene interactions with many intervening genes.

### Selective enhancer subsets are linked to gene expression programs that underlie region-specific development and function

Harnessing region-specific variation in transcription, we next quantified count data at individual TAPEs and used DESeq2 to identify TAPEs that exhibited region-selective expression patterns. This analysis yielded 390 cortical, 776 hippocampal, and 898 striatal putative enhancers (**Fig. 2A-B**; **Supplementary Data Table 2**). Transcription factor binding motif enrichment analysis at TAPES and gene ontology term analysis at predicted TAPE target genes indicated that region-selective TAPEs play integral roles in several region-specific processes (**Fig. 2C**; **Supplementary Data Table 3**). Hippocampus-selective TAPEs were correlated with genes linked to cellular and nervous system development and cell-cell signaling, and they display an enrichment of NEUROG2 (Neurogenin2) binding motifs, a transcription factor crucial for dentate gyrus development (Chen, Lepier, Berninger, Tolkovsky, & Herbert, 2012; Galichet, Guillemot, & Parras, 2008). Similarly, cortex-selective TAPEs correlated with genes implicated in forebrain and CNS development, neuron axonogenesis, and actin polymerization, and exhibit enrichment in binding motifs for SOX5, a transcription factor that regulates neuronal migration and differentiation in the neocortex (Kwan et al., 2008). Likewise, striatum-selective TAPEs correlated with genes important for several amino acid metabolism and modification pathways as well as mitochondrial functions.These TAPEs are enriched in motifs for the transcription factor ISL1, which is required for differentiation of striatonigral pathway projection neurons (Ehrman et al., 2013). Not surprisingly, genes corresponding to these TAPEs also demonstrated region-selective expression, as seen in the examples *Kcnf1, Prox1*, and *Mn1* for cortex, hippocampus, and striatum, respectively (**Fig. 2D-F**). These observations were confirmed by *in situ* hybridization images obtained from the Allen Brain Mouse Atlas.

**Figure 2.**
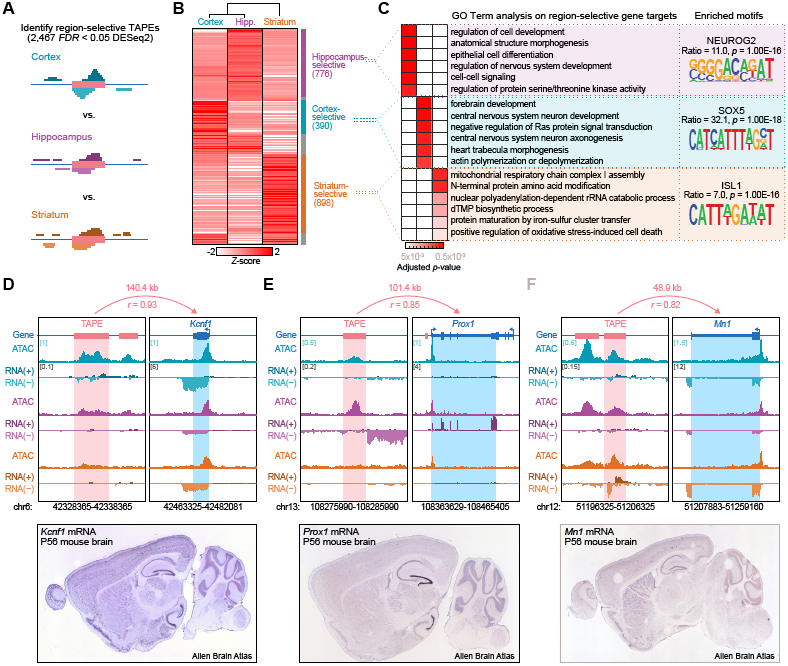
Identification of brain region-selective enhancers and eRNAs in primary neuronal culture systems. **A**, Illustration of DESeq2-based identification of enhancers selective for cortex, hippocampus, and striatum. **B**, Heatmap indicating transcription levels at region-selective TAPEs revealed 776, 390, and 898 TAPEs selective for hippocampus, cortex, and striatum, respectively. **C**, Heatmap showing adjusted *p*-values of the six most enriched Gene Ontology Term groups for genes corresponding to region-selective TAPEs (left). HOMER analysis of transcription factor binding motifs shows an enrichment of characteristic transcription factors within region-selective TAPEs (representative examples on right, complete list in Supplementary Data Table 3). **D-F**, Genome browser tracks showing ATAC-seq and total RNA-seq signal at example loci separated by the three brain regions of interest (top). Region-selective TAPEs are represented by *Kcnf1* for cortex, *Prox1* for hippocampus, and *Mn1* for striatum. Region-selective expression of the respective target genes is also evident in *in situ* hybridization images from the adult mouse brain (bottom, Image credit: Allen Brain Atlas, Allen Institute).

### Activity-dependent enhancer RNA precedes and predicts mRNA induction

To determine whether eRNAs are correlated with activity-dependent alterations in protein-coding genes, we examined RNA transcription from TAPEs following neuronal depolarization with 10 mM potassium chloride (KCl) for 1 hr (**Fig. 3A**). Globally, we identified 96 activity-regulated TAPEs that were significantly altered by KCl treatment in at least one cell type, with 20 selective for cortex, 22 for hippocampus, and 66 for striatum (**Fig. 3B**). Activity-regulated TAPEs demonstrated either increased or decreased transcription in response to neuronal depolarization, here termed upor downregulated TAPEs. As expected, we found that mRNA expression of predicted target genes correlated with TAPE transcription (**Fig. 3C**). However, surprisingly we found that chromatin accessibility (ATAC-seq signal) at either promoter or TAPE regions was not predictive of gene expression levels (**Fig. 3C**), but often shifted in the opposite direction of TAPE/gene RNA estimates. While the decreased TAPE ATAC signal could be specific to this time point, it is worth noting that enhancer transcription in this case provides better predictions of basal and stimulus-dependent gene expression than chromatin accessibility.

**Figure 3.**
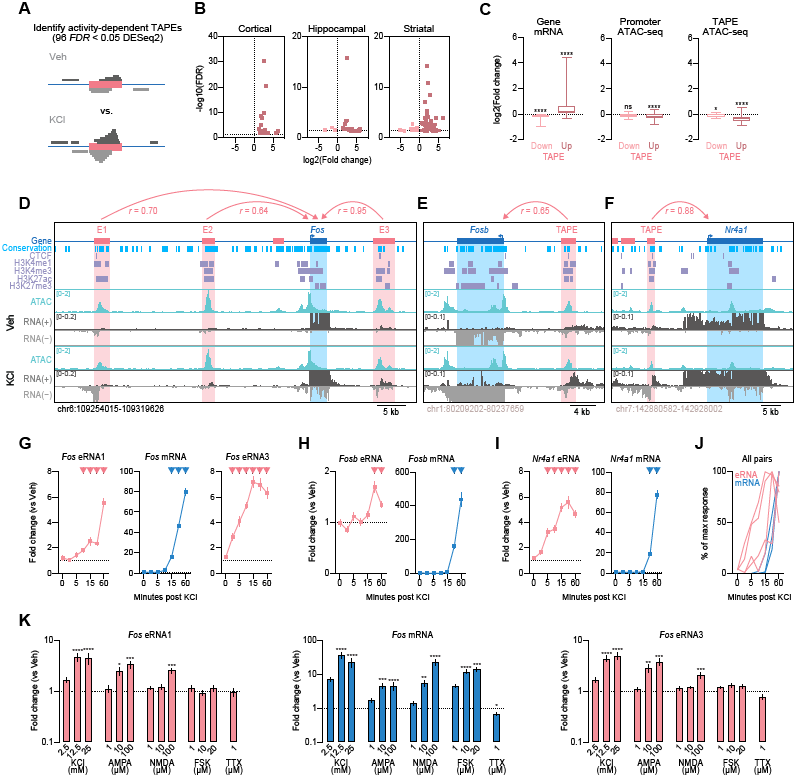
Activity dependence of enhancers and eRNAs. **A**, Illustration of activity-dependent TAPE identification. **B**, Volcano plots showing differentially expressed TAPEs in primary neuronal cultures generated from rat cortex, hippocampus, and striatum. **C**, mRNA levels from TAPE-linked genes are predicted by the direction of eRNA changes at upregulated and downregulated TAPEs (Wilcoxon signed rank test with theoretical median = 0 for upregulated TAPEs, *n* = 77, *p* < 0.0001, and downregulated TAPEs *n* = 15, *p* < 0.0001). Chromatin accessibility decreases at gene promoters that correspond to upregulated TAPEs (Wilcoxon signed rank test with theoretical median = 0 for upregulated TAPEs, *n* = 78, *p* < 0.0001, and downregulated TAPEs, *n* = 15, *p* < 0.0730). ATAC-seq signal also decreases at both up- and downregulated TAPEs (Wilcoxon signed rank test with theoretical median = 0 for upregulated TAPEs, *n* = 78, *p* < 0.0001, and downregulated TAPEs, *n* = 15, *p* < 0.0103). **D-F**, Genomic locus of *Fos, Fosb*, and *Nr4a1* genes and their surrounding enhancer regions. **G-I**, Time course experiments following neuronal depolarization with 25 mM KCl revealed that eRNAs are induced prior (*Fos* eRNA1, eRNA3, and *Nr4a1* eRNA) or at the same time (*Fosb* eRNA) as mRNAs are induced (two-way ANOVA for *Fos* eRNA1, *F*(6,153) = 41.81, *p* < 0.0001, *Fos* eRNA3 *F*(6,154) = 37.87, *p* < 0.0001, *Fos* mRNA, *F*(6,154) = 456, *p* < 0.0001, *Nr4a1* eRNA, *F*(6,154) = 31.4, *p* < 0.0001, *Nr4a1* mRNA, *F*(6,154) = 311.3, *p* < 0.0001, *Fosb* eRNA, *F*(6,154) = 8.341, *p* < 0.0001, *Fosb* mRNA, *F*(6,154) = 98.34, *p* < 0.0001, Sidak’s post hoc test for multiple comparisons). Inverted triangles represent *p* < 0.05 as compared to vehicle treated controls. **J**, Summary of KCl time course experiments plotted as percentage of maximal response. **K**, RT-qPCR analysis of eRNA and mRNA expression in response to 1 hr treatment with KCl, AMPA, and NMDA reveals activity-dependent induction of *Fos* eRNA1 and eRNA3, while FSK and TTX treatment had no effects on *Fos* eRNA expression (Kurskal-Wallis test for eRNA1 KCl *F*(3,32) = 25.04, *p* < 0.0001, AMPA *F*(3,32) = 20.81, *p* = 0.0001, NMDA *F*(3,32) = 17.79, *p* = 0.0005, FSK *F*(3,32) = 1.967, *p* = 0.5793, eRNA3 KCl *F*(3,32) = 26.52, *p* < 0.0001, AMPA *F*(3,32) = 26.11, *p* < 0.0001, NMDA *F*(3,32) = 15.66, *p* = 0.0013, FSK *F*(3,32) = 5.961, *p* = 0.1135, and mRNA KCl *F*(3,32) = 28.26, *p* < 0.0001, AMPA *F*(3,32) = 29.79, *p* < 0.0001, NMDA *F*(3,32) = 29.79, *p* < 0.0005, FSK *F*(3,32) = 30.28, *p* < 0.0001, with Dunn’s post hoc test for multiple comparisons, and unpaired t-test for eRNA1 TTX *t*(14) = 0.1740, *p* = 0.8644, eRNA3 TTX *t*(14) = 1.461, *p* = 0.166, and mRNA TTX *t*(14) = 2.346, *p* = 0.0342). Data expressed as mean ± s.e.m. Multiple comparisons, \**p* < 0.05, \*\**p* < 0.01, *** *p* <0.001, \*\*\*\**p* < 0.0001.

**Figure 3D-F** show ATAC-seq and RNA-seq results from three representative IEGs (*Fos, Fosb*, and *Nr4a1*) that are significantly induced by KCl depolarization. Each of these genes displayed distal activity-regulated TAPEs, including at least three distinct enhancers near the *Fos* gene. The locations of these enhancers are consistent with locations of enhancer elements in other species relative to the *Fos* gene (Joo et al., 2016; Kim et al., 2010) and map to DNA sequences that are enriched for histone modifications associated with active enhancers (H3K4me1 and H3K27ac). Further, each of these elements undergoes bidirectional transcription to yield strand-specific eRNAs. Using ChIP-PCR and RT-qPCR, we also demonstrate that RNAP2 is recruited to the most distal *Fos* enhancer (here termed E1) after neuronal depolarization, and that transcription from this enhancer requires RNAP2 (**Fig. S1**).

To further explore this TAPE-gene relationship, we performed a KCl stimulation time-course experiment in which cultured neurons were depolarized with 25mM KCl, and RNA was isolated from neurons at multiple time points (0, 3.75, 5, 7.5, 15, 30, and 60 min) after treatment. Here, we focused on enhancer RNAs transcribed from the two most distal and most conserved *Fos* enhancers (upstream enhancer-1 and downstream enhancer-3) as eRNAs transcribed from enhancer-2 showed the weakest correlation with *Fos* mRNA in RNA-seq datasets (**Fig. 3D**). RT-qPCR using transcript-specific primers at the *Fos* locus revealed that following KCl depolarization, *Fos* eRNA1 and eRNA3 are significantly upregulated within 7.5 min, whereas *Fos* mRNA is not significantly upregulated until 15 min after stimulation (**Fig. 3G**). We observed similar patterns at *Fosb* and *Nr4a1* loci, indicating that for many IEGs, eRNA induction precedes mRNA induction in response to neuronal depolarization (**Fig. 3H-I**). While we included earlier time points than previous studies to capture the window in which eRNA and mRNA are first induced, our data corroborate the described dynamics of eRNA transcription (Schaukowitch et al., 2014; Arner et al., 2015). Further, our results show that eRNA transcription in response to activity is rapid and follows distinct temporal profiles as compared to mRNA from linked genes (**Fig. 3J**).

To determine whether eRNAs are sensitive to other forms of neuronal and synaptic activation or inactivation, we treated cortical neurons with a variety of pharmacological compounds, including KCl, specific glutamate receptor agonists (AMPA and NMDA), the adenylyl cyclase activator Forskolin (FSK), or the sodium channel blocker tetrodotoxin (TTX) at 11 days *in vitro* (DIV). We found increased transcription of *Fos* eRNA1 and eRNA3 in response to KCl, AMPA, and NMDA in a dose-dependent fashion (**Fig. 3K**). Interestingly, while FSK treatment resulted in strong *Fos* mRNA induction, it did not affect *Fos* eRNA levels. Likewise, only mRNA levels were reduced by TTX. Together, these results suggest that *Fos* eRNA levels are modulated by neuronal activity states in a similar but distinct fashion compared to mRNA levels.

To gain insight into the spatial distribution of eRNAs and their response to stimulation, we performed single molecule fluorescent in situ hybridization (smFISH), a technique that allows visualization of individual eRNA and mRNA transcripts on a single-cell level. Our data confirmed a KCl-mediated induction of *Fos* eRNA1 and revealed a correlation of *Fos* eRNA1 and *Fos* mRNA transcript numbers (**Fig. S2**). These results suggest that eRNAs contribute to transcriptional regulation of their target genes not only on a cell population level, but also on a single-cell level, and that enhancer transcription in single cells may explain at least part of the variability in expression from linked genes.

### Functional validation of enhancer-gene pairs using CRISPR activation tools

To verify predicted enhancer-gene pairs, we sought to determine whether transcriptional activation at selected candidate enhancers was sufficient to induce mRNA at linked genes. To test this, we employed a CRISPR-dCas9 activation (CRISPRa) system in which dCas9 is fused to a strong transcriptional activator (such as VPR or VP64), enabling selective activation of targeted genomic sites (**Fig. 4A-B, Fig. S3**; Savell et al., 2018; K. Li et al., 2020). We designed CRISPR single guide RNAs (sgRNAs) to target the CRISPRa system to transcriptionally active enhancer loci near the *Fos, Fosb*, and *Nr4a1* genes, as well as sgRNAs targeting proximal promoters to drive mRNA transcription directly (**Fig. 4C-D, Fig. S3, Fig S4**). We chose *Fos, Fosb*, and *Nr4a1* based on their dynamic and activity-dependent nature, which makes them promising targets to study the mechanistic interactions between eRNAs and enhancer function in neurons. As a non-targeting negative control, we employed a sgRNA for *lacZ*, a bacterial gene that is not present in eukaryotes.

**Figure 4.**
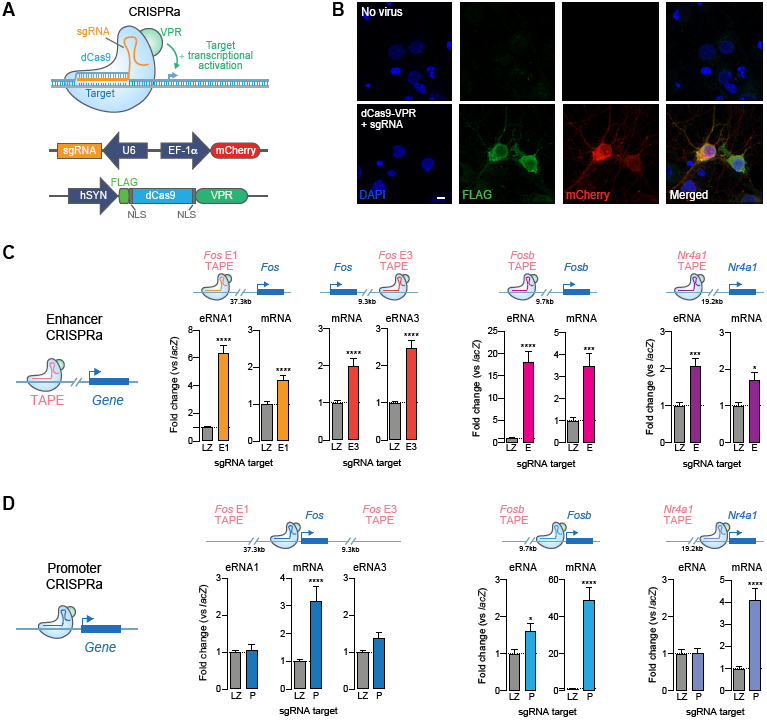
Transcriptional activation at enhancers is sufficient to induce linked genes. **A**, Illustration of CRISPR activation (CRISPRa) strategy for site-specific targeting of the transcriptional activator VPR. **B**, Immunocytochemistry on DIV 11 cortical neurons. Top, no virus control. Bottom, neurons co-transduced with lentiviruses expressing dCas9-VPR (marked by FLAG) and a custom sgRNA (mCherry reporter). Scale bar = 5 μm. **C**, CRISPRa targeting to distal enhancers near *Fos, Fosb*, and *Nr4a1* loci activates eRNA and mRNA from linked genes. Gene expression differences were measured with RT-qPCR (*Fos, n* = 18; *Fosb* and *Nr4a1 n* = 9 per group; two-tailed Mann-Whitney test for all comparisons as compared to non-targeting *lacZ* sgRNA control). **D**, CRISPRa at *Fos, Fosb*, and *Nr4a1* promoters increases mRNA but does not consistently increase eRNA (*Fos, n* = 18; *Fosb* and *Nr4a1 n* = 9 per group; two-tailed Mann-Whitney test for all comparisons as compared to non-targeting *lacZ* sgRNA control). Data expressed as mean ± s.e.m. Multiple comparisons, \**p* < 0.05, \*\**p* < 0.01, \*\*\**p* < 0.001, \*\*\*\**p* < 0.0001.

At DIV 4-5, cortical cultured neurons were transduced with separate lentiviruses expressing dCas9-VPR and sgRNA constructs. On DIV 11, we confirmed transgene expression (indicated by mCherry reporter for sgRNA constructs and FLAG immunocytochemistry for the VPR construct; **Fig. 4B**) and extracted RNA for RT-qPCR. At all four candidate eRNA-mRNA pairs, CRISPRa-mediated transcriptional activation of enhancers not only increased eRNA expression but also significantly induced corresponding mRNA levels (**Fig. 4C**). In contrast, dCas9-VPR targeting to gene promoters specifically increased target mRNA at all candidate genes but did not alter eRNA levels at three out of four candidate pairs (**Fig. 4D**). For example, activation of distinct enhancers either upstream (enhancer-1) or downstream (enhancer-3) of the *Fos* gene produced local eRNA (eRNA1 and eRNA3) induction, but also significantly increased *Fos* mRNA expression. Notably, transcriptional activation of the upstream enhancer did not activate the downstream enhancer, and vice versa (**Fig. S3, Fig. S4**). This strongly indicates that enhancers and eRNAs can be induced by transcriptional activators, that this activation can drive mRNA expression, and that there is little crosstalk between transcriptional activation states at enhancers. Interestingly, dual activation of both enhancers with multiplexed sgRNAs targeting *Fos* enhancer-1 and enhancer-3 had additive effects and stronger mRNA induction compared to individual enhancer activation (**Fig. S4**). Given that enhancers can interact with promoters in enhancer-promoter loops, it is possible that transcriptional activators are close enough to act simultaneously on enhancers and promoters. However, we observed little or no effect on eRNA expression when we targeted gene promoters to drive mRNA expression (**Fig. 4D, Fig. S4**), suggesting that enhancer regulation of linked mRNA is a unidirectional phenomenon. Moreover, we did not observe any effects of enhancer activation on non-targeted eRNAs or mRNAs, supporting the site-specificity of observed CRISPRa effects (**Fig. S4**).

To determine whether these results translate to non-neuronal cell types that were not used to generate enhancer-gene pair predictions, we repeated selected experiments in C6 cells, a rat glioma dividing cell line (**Fig. S3**). As in neurons, we found that recruitment of transcriptional activators (VPR or VP64) to selected *Fos* enhancers not only induced transcription at enhancers but also upregulated *Fos* mRNA. In contrast, *Fos* promoter targeting increased mRNA levels without altering eRNA levels. Together, these findings imply that enhancers can be activated in a site-specific manner and that observed increases in mRNA are due to enhancer activation and potentially increased eRNA levels. Further these results validate enhancer-gene pair predictions based on eRNA transcription abundance.

### Enhancer RNAs are necessary and sufficient for induction of mRNA

To further interrogate the functional role of eRNAs, we explored the effect of eRNA localization on the expression of linked genes. To do so, we employed CRISPR-Display (Shechner et al., 2015), a novel CRISPR approach that allowed us to tether a specific accessory RNA (acRNA) sequence to chosen target sites in the genome and investigate local effects, as compared to global over-expression approaches. Given eRNAs from *Fos* enhancers showed high sequence conservation and robust effects on mRNA expression in neurons as well as in C6 cells, we designed Display acRNA sequences based on conserved regions within the enhancer elements from this gene. We packaged dCas9 along with either sgRNA-eRNA (eRNA-tethering Display construct) or sgRNA-alone (no-acRNA control construct) cassettes into a single plasmid expression vector (**Fig. 5A**). Constructs containing either *Fos* enhancer-1 sgRNA or a non-targeting *lacZ* control were nucleofected into C6 cells, followed by RT-qPCR after a 16 hr incubation period. Anchoring of an acRNA sequence based on *Fos* eRNA1 in close proximity to its parent enhancer (*Fos* enhancer-1) resulted in increased *Fos* mRNA levels compared to the dCas-only control (**Fig. 5B**). Importantly, overexpression of the eRNA1 sequence without enhancer targeting did not affect *Fos* mRNA expression, indicating that the effects of this eRNA are location-dependent (**Fig. 5B, left**). To determine whether the length of the acRNA contributes to the observed effects, we constructed eRNA-tethering CRISPR-Display plasmids with increasing acRNA lengths of 150, 300, and 450 nucleotides (nt; **Fig. 5A, bottom**). Intriguingly, RNA length did not further increase the effect of CRISPR-Display targeting on mRNA expression (**Fig. 5B, right**), suggesting that eRNA-mediated increases in mRNA expression are not directly proportional to the size of eRNA delivered. Since the effects of eRNA targeting appeared to be location-dependent, we next sought to determine their location specificity. We targeted enhancer-1, -3, and a non-regulatory control region between enhancer-1 and the promoter with Display constructs tethering 150 nt long sequences of eRNA1, 3, or a control RNA based on the targeted control region (**Fig. 5A, bottom**). These experiments revealed that only *Fos* eRNA1 tethered to its own enhancer induced mRNA (**Fig. 5C, middle**). Importantly, no significant effects were observed when other RNAs were tethered to enhancer-1 (**Fig. 5C, middle**), nor when eRNA1 was tethered to either enhancer-3 or the control locus (**Fig. 5C, left and right**). These results suggest that the observed effects are specific to eRNA1 when localized to its origin enhancer. Interestingly, eRNA3 did not produce the same effects on mRNA expression as eRNA1 (**Fig. 5C, right**). This could either be due to technical differences between the acRNA designs or hint towards different functional roles between the two eRNAs. While none of the acRNAs contain specific binding motifs that the authors are aware of, it is possible that the chosen 150 nt of eRNA3 does not carry the required properties to be functional in this assay and that a different region of the eRNA3 sequence would show similar results to the eRNA1 acRNA. More importantly, these experiments provide novel evidence that *Fos* eRNA1 acts locally and is sufficient to induce the *Fos* gene.

**Figure 5.**
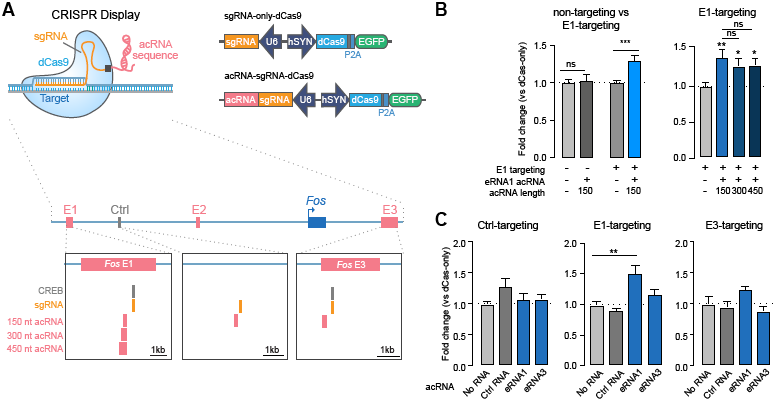
*Fos* eRNA1 is sufficient for *Fos* mRNA expression. **A**, Illustration of Display plasmids (top), and CREB binding motifs, sgRNA target sites, and eRNA regions chosen for CRISPR-Display constructs (acRNA1-3) (bottom). **B**, RT-qPCR analysis reveals that while the *lacZ*-targeting 150 nt eRNA1 acRNA does not affect *Fos* mRNA (*n* = 9 per group, two-tailed Mann-Whitney test, *U* = 36, *p* = 0.7304), targeting eRNA1 to *Fos* enhancer-1 results in increased *Fos* mRNA expression (*n* = 21 per group, two-tailed Mann-Whitney test, *U* = 77, *p* = 0.0002, graph contains data from 12 replicates (from 4 experiments) shown in the right graph and 9 additional replicates (from 2 experiments)). Constructs with increasing acRNA lengths (150, 300, and 450 nt) did not result in stronger *Fos* mRNA induction (one-way ANOVA *F*(3,44) = 3.791, *p* = 0.0167, Tukey’s post hoc test for multiple comparison). **C**, Tethering 150 nt acRNAs based on eRNA1, eRNA3, and a control RNA revealed that only eRNA1 tethering to its own enhancer induced *Fos* mRNA expression (one-way ANOVA for control-targeting *F*(3,19) = 3.191, *p* = 0.0472, and Kurskal-Wallis test for E1-targeting *F*(3,44) = 23.27, *p* < 0.0001, E3 targeting *F*(3,40) = 5.183, *p* = 0.1589). Data expressed as mean ± s.e.m. Multiple comparisons, \**p* < 0.05, \*\**p* < 0.01, \*\*\**p* < 0.001, \*\*\*\**p* < 0.0001.

Based on the CRISPR-Display results, we next sought to address the functional requirement of eRNAs in activity-dependent gene transcription in cortical neurons. We employed an anti-sense oligonucleotide (ASO) strategy to directly target eRNA1 while leaving mRNA and other enhancer functions unperturbed. Rat primary cortical cultures were treated with sequence-specific eRNA1 ASOs for 24 hrs prior to RNA harvesting followed by RT-qPCR (**Fig. 6A**). ASOs targeted to *Fos* eRNA1 induced a robust decrease in eRNA1 expression but did not alter expression of eRNAs from other *Fos* gene enhancers (reinforcing the concept of functional independence of *Fos* eRNAs). Notably, *Fos* eRNA1 ASOs also produced a significant decrease in *Fos* mRNA levels, both at baseline and following neuronal depolarization with KCl (**Fig. 6A, C**). These results suggest that *Fos* eRNA1 is not only required for normal expression from the *Fos* gene, but also for neuronal activity-dependent expression of this IEG. In contrast, we found that knockdown of *Fos* mRNA with an ASO targeted to the mRNA had no effect on eRNA synthesis from any enhancer, further supporting a unidirectional model of eRNA function (**Fig. 6B**). Overall, these findings demonstrate that altering the levels of a single eRNA is sufficient to modulate gene expression.

**Figure 6.**
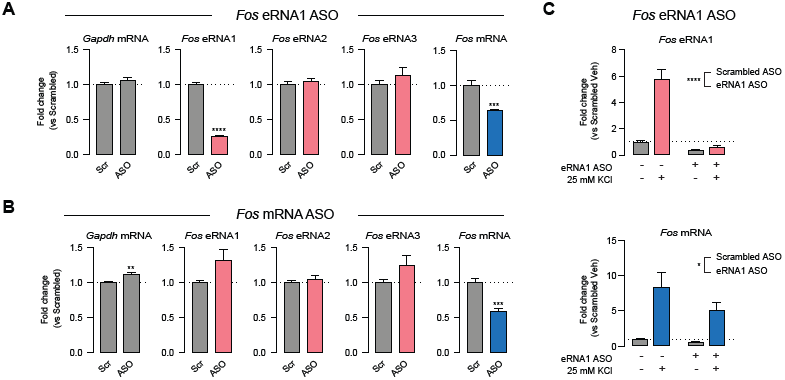
*Fos* eRNA1 is necessary for *Fos* mRNA expression in neurons. **A**, Anti-sense oligonucleotide (ASO) targeting of *Fos* eRNA1 for 24 hrs decreased both eRNA1 and *Fos* mRNA (unpaired t-test *t*(10) = 20.69, *p* < 0.0001 and *t*(10) = 5.739, *p* = 0.0002), but did not alter eRNA levels from other *Fos* enhancers (unpaired t-test for *Gapdh* mRNA *t*(10) = 0.9696, *p* = 0.3551; eRNA2 *t*(10) = 0.8608, *p* = 0.4095; eRNA3 *t*(10) = 1.014, *p* = 0.3346). **B**, *Fos* mRNA targeting ASOs decreased *Fos* mRNA (*t*(10) = 5.198, *p* = 0.0004) with no significant effect on eRNA levels (unpaired t-test for *Gapdh* mRNA *t*(10) = 3.744, *p* = 0.0038; *e*RNA1 *t*(10) = 2.056, *p* = 0.0668; eRNA2 *t*(10) = 0.6508, *p* = 0.5298; eRNA3 *t*(10) = 1.679, *p* = 0.124). **C**, *Fos* eRNA1 ASO pretreatment for 24 hrs prior to 1 hr Veh treatment or KCl stimulation reduced induction of eRNA1 (top) and mRNA (bottom) when compared to a scrambled ASO control (*n* = 9 per group, two-way ANOVA for eRNA1 *F*(1,32) = 154.4 *p* < 0.0001, for mRNA *F*(1,32) = 5.267 *p* = 0.0284). Data expressed as mean ± s.e.m. Multiple comparisons, \**p* < 0.05, \*\**p* < 0.01, \*\*\**p* < 0.001, \*\*\*\**p* < 0.0001.

### Enhancer RNAs interact with but do not depend on chromatin remodelers

The mechanisms by which eRNAs can regulate proximal mRNA transcription remain poorly understood, and while there is evidence for specific eRNA-protein interactions in the literature, no general mechanism has been identified. eRNAs have been demonstrated to be involved in the release of transcriptional repressors, enhancer looping, and epigenetic modifications (Bose et al., 2017; Hsieh et al., 2014; Li et al., 2013; Rahnamoun et al., 2018; Schaukowitch et al., 2014). One possible role of eRNAs lies in the regulation of dynamic chromatin reorganization. Enhancer activation is often accompanied by the recruitment of CBP, CREB, MEF2, NPAS4 and FOS proteins to enhancers near activity-regulated genes (e.g., *Fos, Rgs2*, and *Nr4a2*; Kim et al., 2010). To interrogate the relationship between eRNAs and these enhancer-binding epigenetic modifiers, we focused CBP, which is recruited to enhancers upon activation and has been shown to bind eRNAs (Bose et al., 2017).

First, to determine whether *Fos* eRNA and mRNA expression is regulated by CBP levels, we designed a custom shRNA expression vector targeting *Crebbp* mRNA (which codes for CBP protein). Specific shRNA sequences targeting *Crebbp* mRNA or a scrambled control sequence were cloned into a lentivirus-compatible expression vector and used for lentiviral packaging. At DIV 4-5, cortical cultured neurons were transduced with either control or *Crebbp* shRNA. On DIV 11, we confirmed transgene expression (indicated by mCherry reporter) and extracted RNA for RT-qPCR. Intriguingly, *Crebbp* knockdown reduced *Fos* mRNA by ∼ 40%. However, we observed no reduction in eRNA1 or eRNA3 levels (**Fig. 7A**). Even though CBP has been shown to interact with enhancers, our data indicate that *Crebbp* is crucial for *Fos* mRNA induction but is not required for eRNA transcription.

**Figure 7.**
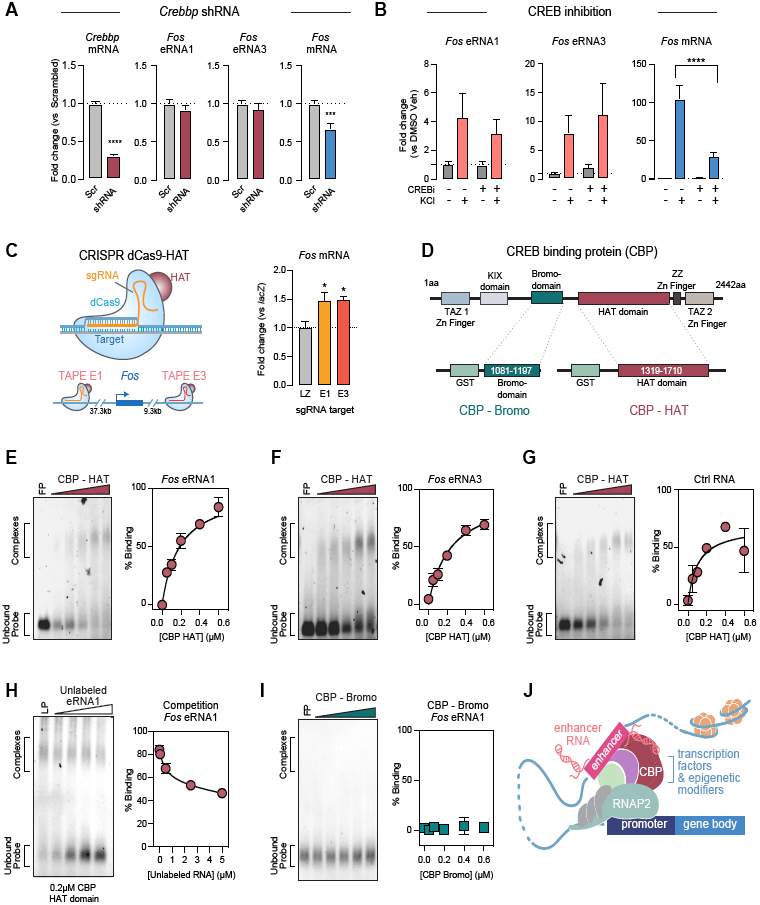
*Fos* eRNA is transcribed independently of but interacts with the histone acetyltransferase CBP. **A**, shRNA-mediated knockdown of *Crebbp* mRNA resulted in deceased *Fos* mRNA but not eRNA expression (*n* = 18 per group, Mann-Whitney for *Crebbp* mRNA *U* = 0 *p* < 0.0001, *Fos* mRNA *U* = 55 *p* = 0.0004, eRNA1 *U* = 140 *p* = 0.5010, eRNA3 *U* = 132 *p* = 0.3550). **B**, CREB inhibition (666-15; 1µM) blunted the KCl response of *Fos* mRNA but not eRNA (*n* = 6 per group, two-way ANOVA for mRNA *F*(1,20) = 37.79 *p* < 0.0001, eRNA1 *F*(1,20) = 0.2461 *p* = 0.8769, eRNA3 *F*(1,20) = 0.1592 *p* = 0.6941, with Tukey’s post hoc test for multiple comparisons). **C**, CRISPR dCas-HAT targeting in C6 cells, in which dCas9 carrying a histone acetyltransferase domain is expressed with sgRNAs to target selected enhancers (left) induced *Fos* mRNA transcription (right, *n* = 8-9 per group, one-way ANOVA *F*(2,23) = 6.151 *p* = 0.0072). **D**, Illustration of CREB-binding protein (CBP) domains (top), and recombinant glutathione-S-transferase (GST) tag-containing CBP-histone acetyltransferase (HAT) domain and CBP-bromodomain (Bromo) used in mobility shift assays. **E-G**, RNA electrophoretic mobility shift assay (REMSA) with escalating concentrations of recombinant protein (0 = free probe (FP), 0.05-0.6 µM) reveal complete binding of synthetic *Fos* eRNA1 (151bp, 50 nM), eRNA3 (153bp, 50 nM), and control RNA (150 bp, 50 nM) to CBP-HAT. **H**, Unlabeled eRNA1 competes for CBP-HAT binding with labeled eRNA1 (LP) in competition assay. **I**, No binding of eRNA1 to CBP-Bromo domain was observed. For all REMSA experiments, *n* = 6 per group. **J**, Model of enhancer function at promoters with associated eRNAs interacting with CBP. Data expressed as mean ± s.e.m. Multiple comparisons, \**p* < 0.05, \*\**p* < 0.01, \*\*\**p* < 0.001, \*\*\*\**p* < 0.0001.

We next tested whether CREB-CBP interactions were critical for *Fos* eRNA and mRNA expression using a selective small molecule compound (666-15) that blocks CREB-mediated gene transcription. DIV 11 cortical neurons were treated with 666-15 and KCl for 1hr followed by RNA harvesting and RT-qPCR. CREB inhibition significantly blunted the *Fos* mRNA response to KCl (**Fig. 7B**). However, CREB inhibition did not alter *Fos* eRNA1 or eRNA3 expression either at baseline or in response to KCl, demonstrating that eRNA expression does not require CREB-CBP interactions.

Since CBP is not required for *Fos* eRNA transcription, we next investigated the role of CBP in enhancer function. First, we tested the effects of CBP’s core histone acetyltransferase (HAT) domain on enhancer function. We paired a sgRNA specific to the *Fos* enhancer-1 locus with a dCas9 fusion containing the core HAT domain from p300, which is nearly identical to the HAT domain from CBP (Hilton et al., 2015). HAT targeting to either *Fos* enhancer in C6 cells elevated *Fos* mRNA to a similar extent as CRISPR-based eRNA1 tethering to the same locus (**Fig. 5B, C, 7C**), suggesting that eRNAs may link enhancer transcription to downstream chromatin remodeling via histone acetylation.

Recent work suggested that eRNAs interact with CBP and modulate the HAT activity of this protein (Bose et al., 2017). However, eRNAs have also been shown to interact with proteins containing a bromodomain (which binds acetylation marks on histones) (Rahnamoun et al., 2018). Intriguingly, CBP contains both a bromodomain and a HAT domain (**Fig. 7D**), implying either function is possible. Furthermore, enhancer acetylation has recently been shown to be a crucial regulatory mechanism of activity-induced dynamic gene expression and transcriptional bursting (Chen et al., 2019). To investigate whether eRNA binding via either of these mechanisms contributes to eRNA function at *Fos* enhancers, we performed RNA electrophoretic mobility shift assays (REMSAs) using ∼ 150nt RNA probe sequence based on *Fos* eRNA1, eRNA3, and a control RNA. REMSA probes were transcribed *in vitro* with fluorescein-labeled bases, and subsequently incubated with recombinant CBP HAT or bromodomains (**Fig. 7E-I**). Strikingly, we observed almost complete binding of *Fos* eRNA1 and the CBP HAT domain at the highest protein concentration (0.6 µM, **Fig. 7E**). While both tested eRNAs bound the CBP HAT domain, it is noteworthy that all tested RNAs, including the control RNA, interacted with the HAT domain. As eRNA1 showed almost complete binding and the strongest effects on mRNA expression in our Display experiments, we used this sequence for control experiments. These experiments demonstrate that the eRNA1-HAT interaction was competitively inhibited by unlabeled eRNA1, suggesting that binding was not mediated by the fluorescent label. Additionally, there was no appreciable binding of eRNA1 to the CBP bromodomain at identical concentrations (**Fig. 7H-I**). This result thus supports recent findings by Bose et al. that various eRNAs can interact with CBP and potentially increase its HAT activity (Bose et al., 2017). Together with our eRNA-tethering results, this data extends the previous findings, suggesting that functional specificity of eRNAs is driven by the location of eRNA transcription rather than their sequence.

Taken together, these results suggest that while eRNAs are not dependent on CREB and CBP, they can facilitate transcriptional induction through direct interaction with the CBP HAT domain. Our findings suggest a model for eRNA function in which eRNAs participate in enhancer-promoter communication where they can interact with epigenetic modifiers such as CBP (**Fig. 7J**).

## DISCUSSION

Distal enhancer elements in DNA enable higher-order chromatin interactions that facilitate gene expression programs and thus contribute to cellular phenotype and function (Heinz et al., 2015; Li et al., 2016; Wang et al., 2011). In the developing brain, the majority of enhancer elements exhibit temporally specific emergence during precise developmental windows, with only ∼15% of enhancers being utilized continually from late embryonic development into adulthood (Gray et al., 2015; Nord et al., 2013). These developmentally regulated enhancers contribute to cell- and tissue-specific gene expression patterns that establish communication within and between brain structures (Frank et al., 2015; Nord et al., 2013; Pattabiraman et al., 2014). Not surprisingly, enhancers utilized in early embryonic brain development possess the highest degree of sequence conservation across species, suggesting that robust evolutionary pressures drive enhancer function (Nord et al., 2013). In the postnatal and mature brain, enhancers continue to play a widespread role in the activity-dependent transcriptional programs that regulate key aspects of neuronal plasticity and function (Gray et al., 2015; Hnisz et al., 2013; Joo et al., 2016; Kim et al., 2010; Malik et al., 2014; Telese et al., 2015; Vermunt et al., 2014, Baizabal et al., 2018; Tyssowski et al., 2018). Repression or deletion of enhancer elements has profound effects on the genes that they control, including complete inactivation (Joo et al., 2016; Kearns et al., 2015; Malik et al., 2014; Telese et al., 2015). Likewise, targeted enhancer activation induces robust upregulation of linked genes, suggesting that enhancers serve as bidirectional regulators of gene activity (Frank et al., 2015; Hilton et al., 2015).

While genome-wide enhancer identification traditionally relies on extensive ChIP-seq, ATAC-seq, and/or HiC-seq datasets to asses location and activity states, our findings are in line with recent work by Wang et al. and emphasize the advantages of transcriptional information as an indicator of local chromatin states (Wang et al., 2020). In this study, the authors developed a machine learning tool to predict chromatin landscapes with nucleosome resolution based exclusively on nascent transcription. Along with this line of work, others have underlined the tight relationship between enhancer transcription and transcription factor activity (Azofeifa et al., 2018), as well as enhancer and promoter function (Mikhaylichenko et al., 2018). Our results demonstrate that eRNA can be a useful measure of enhancer activity state, and often generates a distinct readout from that provided by ATAC-seq signal at the same enhancer. For example, whereas we detected many eRNAs induced by neuronal depolarization, we observed that ATAC-seq signals at these activated enhancers (or the promoters of linked genes) decreased following depolarization. While the molecular origins of this effect are unclear, this finding may suggest that at the timescales examined here, ATAC-seq quantification cannot discriminate between active and poised enhancer states for constitutively poised activity-regulated enhancers. Nevertheless, these results demonstrate that it is possible to resolve activity states of regulatory elements and their corresponding target genes with only a few datasets and suggests that eRNA induction levels can predict mRNA response to stimulation more reliably than changes in chromatin accessibility.

Although it is well accepted that genomic enhancers play critical roles in tuning the spatiotemporal nature of transcription from linked genes, techniques typically used to examine enhancer function (e.g., enhancer deletion (Leighton et al., 1995), Cas9-based mutation (Lopes et al., 2016; Sanjana et al., 2016), or activation/inactivation with dCas9 fusion proteins (Hilton et al., 2015; Thakore et al., 2015; Joo et al., 2016; Liu et al., 2017; Liu et al., 2016; K. Li et al., 2019a)) interfere with both the genomic locus and eRNAs transcribed from that locus. Therefore, these approaches cannot dissociate the effects of enhancer function and eRNA function. To address this problem, we first implemented genome-wide transcriptional profiling in our enhancer identification pipeline to gain a better understanding of the relationship between enhancer transcription, enhancer activity, and downstream transcriptional regulation. Moreover, we took two different approaches that directly target eRNAs in order to examine their function separately from enhancer function. First, we used a novel CRISPR-Display approach to target *Fos* eRNAs to their own enhancer. These results demonstrate that *Fos* eRNA1 is sufficient to induce *Fos* mRNA and provide novel evidence for a location-dependent functional role of eRNA (**Fig. 5**). Furthermore, our data shows that while *Fos* eRNA expression is independent of CREB and CBP function, they can interact with CBP through direct binding to the HAT domain. Likewise, CRISPR-dCas9 mediated recruitment of a HAT domain recapitulated the effects of eRNA-tethering at the same *Fos* enhancer loci (**Fig. 7**). Secondly, we employed stable, cell-penetrating ASOs to target eRNA for degradation. These results suggest that eRNA is necessary for normal expression of *Fos* mRNA, both under basal conditions and after neuronal depolarization. (**Fig. 6**).

Overall, these results agree with a previous report demonstrating that eRNAs transcribed from activity-dependent enhancers are necessary for induction of mRNA from linked genes (Schaukowitch et al., 2014). This report utilized lentiviral shRNA knockdown approaches to directly target activity-induced eRNAs near *Arc* and *Gadd45b* genes, and followed this knockdown with KCl depolarization to induce mRNAs. Targeted shRNA knockdown of eRNA specifically blocked mRNA induction at these genes but not other IEGs induced by neuronal activation (*Fos, Egr1*). Our results extend these important findings in two ways. First, given that the *Fos* gene exhibits multiple enhancers and activity-dependent eRNAs, we were able to address the functional relationship between eRNAs near the same gene. Our results suggest that while eRNAs do regulate mRNA induction at linked genes, eRNAs are functionally independent of each other. Thus, ASO-mediated knockdown of eRNAs transcribed from the most distal *Fos* enhancer did not downregulate eRNAs transcribed from other enhancers (**Fig. 6**). Secondly, in parallel experiments we were able to target *Fos* mRNA for knockdown using an identical approach. These results demonstrate that the relationship between eRNA and mRNA levels at the same gene is unidirectional – i.e., that mRNA knockdown does not also reduce eRNA levels. This is a critical control at autoregulating IEGs like *Fos*, given that the protein product of this gene is a transcription factor that localizes to enhancers in an AP1 complex with Jun family members (Malik et al., 2014).

Biological roles of lncRNAs are generally linked to their ability to bind functionally active proteins to operate as molecular guides, decoy molecules, scaffolding, or even allosteric modulators (Quinn and Chang, 2016; Rinn and Chang, 2012). In agreement with this concept, a large number of chromatin-associated proteins bind RNA in addition to DNA (Di Ruscio et al., 2013; Hendrickson et al., 2016; Savell et al., 2016), and several well-characterized transcriptional regulators have recently been shown to possess functional interactions with eRNAs (Bose et al., 2017; Hsieh et al., 2014; Lai et al., 2013; Li et al., 2013; Li et al., 2016; Schaukowitch et al., 2014; Sigova et al., 2015). For example, eRNAs have been shown to bind the ubiquitous transcription factor Yin-Yang 1 (YY1) to “trap” YY1 at the enhancer, thus facilitating its action at local YY1 motifs in DNA (Sigova et al., 2015). In this study, a similar CRISPR-dCas9 system was used to tether *Arid1a* RNA in close proximity to an enhancer YY1-binding motif. Our CRISPR-Display experiments build on this work by showing direct changes in target gene expression that are dependent on the target location but not sequence length. Our data confirms and highlights the importance and sufficiency of eRNAs as transcriptional organizers.

Similarly, eRNAs can act as decoy molecules for negative elongation factor (NELF) complexes, which are important regulators of RNAP2 pausing and transcriptional bursting (Schaukowitch et al., 2014). In line with previous findings that eRNAs interact with CBP and stimulate its activity as a HAT at enhancer loci (Bose et al., 2017), we found that *Fos* eRNAs can interact with CBP through direct binding to the HAT domain and that eRNA or HAT recruitment to the enhancer increases mRNA expression. Additionally, our data suggest that the specificity of this interaction is based on the location of eRNA transcription, likely in combination with their short half-life rather than sequence specificity.

While our results do not rule out other eRNA-protein interactions, our findings are consistent with recent observations that histone acetylation plays a key role at enhancers by influencing transcriptional properties of corresponding genes (Chen et al., 2019). This study demonstrated that histone acetylation at enhancers increases transcriptional bursting at linked genes, and that these effects are mediated by BRD4, a bromodomain-containing protein important for RNAP2 phosphorylation and transcription. Our results provide the first evidence that eRNA function is dependent on eRNA location and partially dependent on sequence, but not sequence length. Furutre studies will be required to explore how different factors, such as distance from their target gene, influence enhancer and eRNA function and characterize other eRNA/protein interactions in more detail. This will help to determine whether common regulatory mechanisms dictate expression of different eRNAs targeting the same gene, as we observed some characteristic and functional differences between *Fos* eRNA1 and eRNA3. It will further help to unravel whether a group of eRNAs that regulate the same gene have distinct functional mechanisms. Finally, it will be crucial to understand the interplay of different enhancers and eRNAs and how they orchestrate gene expression programs and potentially fine-tune responses to specific stimuli.

The vast majority of gene variants linked to human health and disease by genome-wide association studies are located in non-coding regions of the genome (Gordon and Lyonnet, 2014; Network and Pathway Analysis Subgroup of Psychiatric Genomics, 2015; Schizophrenia Working Group of the Psychiatric Genomics, 2014; Vermunt et al., 2014), with putative enhancers containing more disease-linked single-nucleotide polymorphisms than all other genetic loci combined (Corradin and Scacheri, 2014). Disease-linked genetic variants could affect enhancer activity either via direct modification of enhancer DNA sequence (e.g., disruption of a transcription factor motif) or by alterations in long-range chromatin interactions between enhancers and gene promoters (reviewed in Carullo & Day, 2019). Indeed, numerous diseases have already been linked to sequence variations in enhancer regions (Gordon and Lyonnet, 2014; Jeong et al., 2008; Spieler et al., 2014; Vermunt et al., 2014; Song et al., 2019), including complex polygenic conditions such as depression (Davidson et al., 2011; Edwards et al., 2012), obesity (Davidson et al., 2011; Voisin et al., 2015), schizophrenia (Eckart et al., 2016; Roussos et al., 2014), bipolar disorder (Eckart et al., 2016), Alzheimer’s disease (P. Li et al., 2019b; Nott et al., 2019), and autism spectrum disorders (Inoue and Inoue, 2016; Yao et al., 2015). This growing link between enhancer activity and brain function strongly highlights the need to better understand the mechanistic interactions that regulate enhancer function at the molecular level, and also suggests that enhancers could be attractive targets for a new generation of disease therapeutics.

## Acknowledgements

We thank Allison Bauman for generating our primary neuronal cell cultures and all current and former Day Lab members for assistance and support. We acknowledge the Civitan International Research Center Cellular Imaging Facility. This work was supported by NIH grants MH114990, DA039650, and DA034681 and the UAB Pittman Scholar Program (JJD), R21DA041878, R01DA036865, and the Allen Distinguished Investigator Award from the Paul G. Allen Frontiers Group (CAG), and the CIRC Emerging Scholar Award (NVNC).

## Author contributions

N.V.N.C., R.A.P.III, R.C.S. and J.J.D. conceived of and performed experiments. N.V.N.C. and J.J.D. wrote the manuscript with help from R.C.S.. R.A.P.III performed RNA-sequencing and ATAC-sequencing experiments and performed genome alignment with custom scripts written by L.I. R.A.P.III performed ATAC-seq peak calling on all sequencing datasets and performed TAPE-gene correlation analyses, HOMER analysis, and wrote methods sections describing these approaches. J.J.D. generated enhancer identification pipeline with assistance from R.A.P.III, enrichment analyses in **Figure 1**, as well as identification of region and activity selective expression. N.V.N.C. performed gene ontology analysis, and experiments for time course and dose-response studies with assistance from S.R. and J.H.. FISH experiments and analysis were performed by N.V.N.C. with assistance from A.S.. VPR and VP64 constructs driven by the hSYN promoter were developed by K.E.S.. Lentivirus production protocols were optimized by K.E.S., S.B., and F.A.S.. CRISPR sgRNAs were designed and cloned by N.V.N.C., R.C.S., K.E.S., and F.A.S.. Nucleofection protocols were optimized by N.V.N.C., K.E.S., and J.S.R.. Lentivirus production, transductions, and nucleofections were performed by N.V.N.C. with help and assistance from K.E.S., F.A.S., S.B., A.S., S.R., and J.H.. ICC and imaging was performed by N.V.N.C. with assistance from A.S.. CRISPR-Display plasmids were designed and built by N.V.N.C. with help from J.S.R. based on a previous version designed and cloned by F.A.S. and N.V.N.C. with help from K.E.S.. CRISPR-Display experiments were performed by N.V.N.C. with assistance from A.S., S.R., and J.H. ASOs were designed by J.J.D. and R.C.S. and experiments were designed and performed by R.C.S. and K.D.B.. R.C.S. wrote the ASO methods section and performed RNAPol2 ChIP and inhibitor experiments. CREB inhibition experiments were performed by N.V.N.C. with assistance from S.R. and J.H.. shRNA sequences were cloned by N.V.N.C. into an empty shRNA construct made by F.A.S., with help from S.B.. Knockdown experiments including virus production were performed by N.V.N.C with assistance from S.R. and J.H.. REMSA experiments including oligo synthesis were designed and carried out by N.V.N.C. with assistance from S.R.. CRISPR dCas9-p300 plasmid and sgRNA cloning vectors, as well as advice for using CRISPR tools, were provided by C.A.G.. J.J.D. supervised all work.

## STAR METHODS

### Cultured neuron experiments

Primary rat neuronal cultures were generated from embryonic day 18 rat cortical, hippocampal, or striatal tissue as described previously (Day et al., 2013; Savell et al., 2016). Briefly, cell culture wells were coated overnight at 37° C with poly-L-lysine (0.05 mg/ml for culture wells supplemented with up to 0.05 mg/ml Laminin) and rinsed with diH2O. Dissected tissues were incubated with papain for 25 min at 37°C. After rinsing in Hank’s Balanced Salt Solution (HBSS), a single cell suspension of the tissue was re-suspended in Neurobasal media (Invitrogen) by trituration through a series of large to small fire-polished Pasteur pipets. Primary neuronal cells were passed through a 100 µM cell strainer, spun and re-suspended in fresh media. Cells were then counted and plated to a density of 125,000 cells per well on 24-well culture plate and 250,000 cells per well on 12-well culture plate with or without glass coverslips (60,000 cells/cm). Cells were grown in Neurobasal media plus B-27 and L-glutamine supplement (complete Neurobasal media) for 11 DIV in a humidified CO2 (5%) incubator at 37° C.

At 4-11 DIV, neuronal cultures were treated as described. For KCl stimulation experiments, KCl (Sigma) was added to complete Neurobasal media, and KCl solution or vehicle (complete Neurobasal media alone) was added for the indicated final concentrations. Cells were incubated with KCl for described time points prior to RNA extraction. For TTX inactivation experiments, cells were treated with 1 µM TTX (Tocris Bioscience) in Neurobasal media for the described time points prior to RNA extraction. S-AMPA, NMDA, and FSK (Sigma) were resuspended in sterile water, diluted in Neurobasal media, and added to cultures for 1 hr at equal volumes (final concentrations of 1 µM, 10 µM, or 100 µM). Same volume of Neurobasal media was added as a vehicle control. For experiments involving RNAP inhibitors, cultures were treated for 4 hrs or 4 hrs followed by a 1 hr, 25 mM KCl stimulation. The RNAP2-dependent transcriptional inhibitor DRB (Sigma) was dissolved to a 20 mM stock solution in 100% cell culture grade DMSO (Sigma) and diluted in Neurobasal media to described experimental concentrations. For CREB inhibitor experiments CREBi (666-15, Torcis) also called 3-(3-Aminopropoxy)-N-[2-[[3- [[(4-chloro-2-hydroxyphenyl)amino]carbonyl]-2-naphthalenyl]oxy]ethyl]-2-naphthalenecarboxamine hydrochloride was dissolved to a 10 mM stock solution in 100% cell culture grade DMSO (Invitrogen) and diluted in Neurobasal media to for a final treatment concentration of 1µM. In experiments involving DRB and 666-15, vehicle-treated cells received equal concentrations of DMSO in Neurobasal media.

For viral transduction, cells were transduced with lentiviruses on DIV 4 or 5 (only viruses with a minimum titer of 1×109 GC/ml were used for target multiplicity of infection (MOIs) of at least 1000). After an 8-16 hr incubation period, virus-containing media was replaced with conditioned media to minimize toxicity. A regular half-media change followed on DIV 8. On DIV 11, transduced cells were imaged and virus expression was verified prior to KCl-treatment and/or RNA extraction. Immunocytochemistry for FLAG was performed as described previously (Savell et al., 2016) with an anti-FLAG antibody (MA1-91878, Thermo Fisher Scientific, RRID AB_1957945). EGFP and mCherry expression was also used to visualize successful transduction using a Nikon TiS inverted epifluorescence microscope.

### RNA extraction and RT-qPCR

Total RNA was extracted (RNAeasy kit, Qiagen) with DNase treatment (RNase free DNAse, Qiagen), and reverse-transcribed (iScript cDNA Synthesis Kit, BioRad). cDNA was subject to RT-qPCR for genes of interest, as described previously (Savell et al., 2016). A list of PCR primer sequences is provided in Supplementary Data Table 4.

### Assay for Transposase-Accessible Chromatin using Sequencing (ATAC-Seq)

Nuclei from rat embryonic cortical, hippocampal, or striatal neurons (50,000/region) were used for ATAC-seq library preparation, following a modified protocol (Buenrostro, Wu, Chang, & Greenleaf, 2015; Corces et al., 2017; Scharer et al., 2016). Briefly, 25 µl of 0.4 M KCl (Sigma) or vehicle (Neurobasal media) was added to cell culture wells to achieve a final concentration of 10 mM KCl and incubated at 37°C for 1 hr. Following treatment, media containing KCl or Vehicle was aspirated, and cells were washed with 1x cold PBS. Then, 600 µl of Lysis Buffer (10 mM Tris-HCl, 10 mM NaCl, 3 mM MgCl2, 0.1% IGEPAL CA-630, Molecular Grade H2O) was added to cell culture wells and incubated for 5 mins on ice. Lysed cells were then transferred to a 1.5 ml Eppendorf tube and centrifuged at 500 g for 10 mins in a swinging bucket centrifuge. Following removal of supernatant containing lysis buffer and cell debris, 500 µl of 1x cold PBS was added to each tube, and nuclei were counted using the Countess II (Life Technologies). The approximate volume needed to achieve 50,000 nuclei was then placed in new Eppendorf tube, and centrifuged at 500 g for 10 mins in a swinging bucket centrifuge. The nuclei pellet was then resuspended in 22.5 µl of tagmentation reaction mix containing Molecular Grade H2O, 2x Tagment DNA Buffer (Illumina), 0.011% Digitonin, 0.1% Tween-20, followed by the addition of 2.5 µl TDE1 (Illumina) and incubated at 37°C for 1 hr. Following tagmentation, libraries were PCR amplified and purified using the Qiagen Minelute PCR purification kit (Qiagen). To remove primer dimers, libraries were bead purified using one round of 1.0X Ampure XP beads with 80% ethanol, followed by a second round of 0.8X Ampure beads with 80% ethanol. Libraries were sequenced (75-bp paired end reads) on the NextSeq500 platform (Illumina).

### ATAC-Seq Mapping and Peak Calling

Paired-end FASTQ files were aligned and trimmed using a custom bioinformatic pipeline initialized in snakemake v5.3.0 (Köster & Rahmann, 2018). Briefly, low-quality bases (Phred <20) and Nextera adapters (5’-CTGTCTCTTATA-3’) were identified and trimmed using FastQC and TrimGalore (v0.4.5), respectively. FASTQ files containing trimmed sequences were then aligned to Rn6 Ensembl genome assembly (v95) to generate binary alignment map (BAM) files with Bowtie2 (v.2.3.4.2) with custom options: ‘--local --very-sensitive-local’, and ‘-X 3000’ (Langmead & Salzberg, 2012). Sequences were then ordered by genomic position using Samtools (v.1.9) (H. Li et al., 2009). Next, BAM files for each brain region were merged to generate three metasamples that were used for downstream data analysis. Peaks for each region were called using MACS2 (Zhang et al., 2008) callpeak with the options - -qvalue 0.00001 - -gsize 2729862089 - -format BAMPE. These options utilize the default behavior of the MACS2 algorithm to ignore duplicates. Within each brain region, peaks closer than 1000bps were merged with BEDtools (Quinlan & Hall, 2010) and peaks less than 146bp, the specific length of DNA wrapped around a single nucleosome, were removed in R. Finally, peaks from each brain region were merged with BEDtools to create an additive peak set containing 192,830 peaks.

### Bulk RNA-Sequencing

Bulk RNA-sequencing (RNA- seq) was carried out at the Heflin Center for Genomic Science Genomics Core Laboratories at the University of Alabama at Birmingham. RNA was extracted, purified (RNeasy, Qiagen), and DNase-treated for three to four biological replicates per brain region and experimental condition. 1 μg of total RNA underwent quality control (Bioanalyzer) and was prepared for directional RNA sequencing using NEBNext reagents (New England Biolabs) according to manufacturer’s recommendations. Specifically, the NEBnext rRNA depletion kit was used to remove ribosomal RNA to include both polyadenylated and non-polyadenylated RNA. RNA-seq libraries underwent sequencing (75 bp paired-end directional reads; ∼12.4-31.1M reads/sample) on an Illumina sequencing platform (NextSeq500).

### Bulk RNA-Seq Data Analysis

Paired-end FASTQ files were uploaded to the University of Alabama at Birmingham’s High-Performance Computer cluster for custom bioinformatics analysis using a pipeline built with snakemake (v5.3.0; Köster & Rahmann, 2018). Read quality was assessed with FastQC and low-quality bases (Phred <20) and Illumina adapters were trimmed with TrimGalore (v0.4.5). Splice-aware alignment to the Rn6 Ensembl genome assembly (v95) was performed with STAR (v2.6.0; Dobin et al., 2013). Binary alignment map (BAM) files were merged and indexed with Samtools (v1.9).

### HOMER Motif Analysis

Region-specific TAPE genomic positions were compiled into three separate BED files. The findMotifsGenome.pl function within the HOMER (v4.11.1) package was used to identify enriched motifs and their corresponding transcription factors within these genomic positions with options -size 1000 -len 8,10,12 -mask -preparse -dumpfasta. As we were interested in the motifs that were specific to each region, a BED file containing peaks from the two other regions was used as background. For example, when identifying enriched motifs for striatal-specific TAPEs, a background file containing genomic positions for cortical- and hippocampal-specific TAPEs. Furthermore, only those de-novo motifs not marked as possible false positives are reported.

### ROC & TAPE Identification

ATAC-seq peaks were used to identify 191,857 ROCs spanning 500 bp up- and down- stream of ATAC-seq peaks. Due to the difficulty in separating intronic enhancers from potential promoters or other elements, the identified ROCs were then filtered for regions that fall >1kb outside of genes curated by Refseq, UCSC, and Ensemble. To account for unannotated or misannotated genes, we filtered for ROCs that don’t overlap contiguously transcribed regions (>100 bp, merging elements closer than 1 kb) or known non-coding RNAs such as miRNA, rRNA, snoRNA, snRNA, and tRNA. These filtering steps provided a list of iROCs. The remaining iROCs were overlaid with RNA-seq data with read cutoffs to select for bidirectionally transcribed loci only to map 28,492 TAPEs. Histone modification peaks from mouse forebrain at postnatal day zero and CTCF binding (ENCODE project datasets obtained from the UCSC Table Browser and transformed from mm10 to Rn6 genome coordinates using Liftover) were quantified at identified TAPEs. Possible TAPE-gene pairs were identified by mapping all gene promoters within 1 Mb upstream or downstream from the center of the TAPE. CPKM values for each TAPE and associated gene were correlated using a Pearson’s correlation in R. TAPE-gene pairs with global correlations of NA were removed as these values were due to the corresponding gene containing count values of 0 for every sample. Removing these pairs left 388,605 potential TAPE- gene pairs. TAPE-gene pairs with correlations greater than 0.5 were deemed high-confidence pairs. These high-confidence pairs were then used to investigate distance and gene position distributions.

### CRISPR-dCas9 and shRNA construct design

To achieve transcriptional activation, lentivirus-compatible plasmids were engineered to express dCas9 fused to VP64 or VPR constructs (Addgene plasmid # 114196 (Savell et al., 2018)). dCAS9-VP64_GFP was a gift from Feng Zhang (Addgene plasmid # 61422 (Konermann et al., 2015)). The pcD- NA-dCas9-p300 Core construct was a gift from Charles Gersbach (Addgene plasmid # 61357 (Hilton et al., 2015)). VP64- and VPR-expressing constructs were co-transduced with sgRNA-containing constructs. A guide RNA scaffold (a gift from Charles Gersbach, Addgene #47108) (Perez-Pinera et al. 2013) was inserted into a lentivirus-compatible backbone, and EF1α-mCherry was inserted for live-cell visualization. A BbsI cut site within the mCherry construct was mutated with a site-directed mutagenesis kit (NEB). Gene-specific gRNAs were designed using an online sgRNA tool, provided by the Zhang Lab at MIT (crispr.mit.edu). To ensure specificity, all CRISPR crRNA sequences were analyzed with the National Center for Biotechnology Information’s (NCBI) Basic Local Alignment Search Tool (BLAST). sgRNAs were designed to target *Fos, Fosb*, and *Nr4a1* enhancers, respectively, as well as the promoter and control regions (a list of the target sequences is provided in Supplementary Data Table 4). crRNA sequences were annealed and ligated into the sgRNA scaffold using the BbsI or BsmBI cut site. For CRISPR-Display, lentiCRISPR v2 from Feng Zhang (Addgene plasmid # 52961 (Sanjana et al., 2014)) was modified and engineered to express dCas9 (instead of Cas9) and GFP under an hSYN promoter, as well as additional restriction sites (Esp3I) for subsequent acRNA insertion using PacI and Xbal. sgRNA, and acRNA sequences of eRNA1, eRNA3 and control RNA were inserted via restriction enzyme cloning using gBlocks for acRNA insertion (cut with Esp3I). As another control, a plasmid lacking the eRNA sequence was targeted to the same genomic sites. To achieve RNA knockdown, shRNA sequences targeting the gene of interest were designed using the Broad

TRC shRNA design tool (http://portals.broadinstitute.org/gpp/public/) according to the Addgene pLKO.1 protocol (https://www.addgene.org/tools/protocols/plko/) and inserted into a lentivirus-compatible shRNA construct (Zipperly et al., 2020). To ensure specificity all shRNA sequences were analyzed with BLAST. All targeting and scrambled control shRNA sequences were annealed and ligated into the shRNA cassette-containing construct using the AgeI and EcoRI cut sites. Plasmids were sequence-verified with Sanger sequencing; final crRNA insertion was verified using PCR.

### Allen Brain Atlas Images

*In situ* hybridization images were obtained from the Allen Mouse Brain Atlas for *Kcnf1* (https://mouse.brain-map.org/experiment/show/68798944), *Mn1* (https://mouse.brain-map.org/gene/show/130518), and *Prox1* (https://mouse.brain-map.org/gene/show/18893).

### C6 Cell Culturing and Nucleofection

C6 cells were obtained from American Type Culture Collection (CCL-107, ATCC, RRID:CVCL_0194) and cultured in F-12k-based medium (2.5% bovine serum, 12% horse serum). At each passage, cells were trypsinized for 1-3 min (0.25% trypsin and 1 mM EDTA in PBS pH 7.4) at room temperature. After each passage remaining cells were processed for nucleofection (2 x106 /group). Cell pellets were resuspended in nucleofection buffer (5 mM KCl, 15 mM MgCl, 15 mM HEPES, 125 mM Na2HPO4/NaH2PO4, 25 mM Mannitol) and electroporated with 3.4 μg plasmid DNA per group. Nucleofector™2b device (Lonza) was used according to the manufacturer’s instruction (C6, high efficiency protocol). Nucleofection groups were diluted with 500 μl media respectively and plated in triplicates in 24-well plates (∼ 666,667 cells/well). Plates underwent a full media change 4-6 hrs after nucleofection and were imaged and frozen for downstream processing after 16 hrs.

### Lentivirus production

Viruses were produced in a sterile environment subject to BSL-2 safety by transfecting HEK- 293T cells with specified CRISPR-dCas9 plasmids, the ps- PAX2 packaging plasmid, and the pCMV-VSV-G envelope plasmid (Addgene 12260 & 8454) with FuGene HD (Promega) for 40-48 hrs as previously described (Savell, Sultan, & Day, 2019). Viruses were purified using filter (0.45 μm) and ultracentrifugation (25,000 rpm, 1 hr 45 min) steps. Viral titer was determined using a qPCR Lentivirus Titration Kit (Lenti-X, qRT-PCR Titration Kit, Takara). For smaller scale virus preparation, each sgRNA plasmid was transfected in a 12-well culture plate as described above. After 40-48 hr, lentiviruses were concentrated with Lenti-X concentrator (Takara), resuspended in sterile PBS, and used immediately. Viruses were stored in sterile PBS at -80°C in single-use aliquots.

### Antisense oligonucleotide (ASO) design and treatment

To manipulate *Fos* mRNA or eRNA levels, we designed 20 bp ASOs that targeted distinct transcripts from the *Fos* gene locus (see Supplementary Data Table 4 for target sequences). ASOs targeting exon 3 of *Fos* mRNA or *Fos* eRNA1 were synthesized with two chemical modifications: an all phosphorothioate backbone and five 2’ O-methyl RNA bases on each end of the oligonucleotide (Integrated DNA Technologies). Primary neuronal cultures were treated with scrambled or *Fos* targeted ASOs (15 μM in buffer EB, for a final concentration of 1.5 μM) and incubated for 24 hrs (basal experiments) or 23 hrs followed by 1 hr neuronal depolarization with 25 mM KCl (or vehicle control). Following ASO treatment, RNA was extracted (Qiagen RNeasy kit) and *Fos* mRNA and eRNA levels were determined using RT-qPCR with custom primers.

### Single Molecule Fluorescent In Situ Hybridisation (sm- FISH)

#### smFISH RNA Probe Design

We designed and ordered Stellaris® FISH probe sets for *Gapdh* mRNA, *Fos* eRNA1, *Fos* eRNA3 and *Fos* mRNA carrying a fluorophore (Quasar® 570 for eRNA1 and *Gapdh* mRNA probes, Quasar® 670 for both *Fos* and *Gapdh* mRNA probes). We preferred probes of 20-mer oligonucleotides. Multiple probes per set targeting the same RNA molecule were designed for an adequate signal to background ratio and to optimize signal strength. Target sequences of each probe set are provided in Supplementary Data Table 4).

#### Sample Preparation and Hybridization

Day 1: Primary neuronal cultures (∼250,000 neurons per coverslip/well) were KCl- or vehicle-treated for 1 hr on DIV 11. After treatment cells were cross-linked with 3.7% formaldehyde (paraformaldehyde in 1X PBS) for 10 min at room temperature (21°C) on a rocking platform. Wells were washed twice with PBS and permeabilized in 70% ethanol for at least 3 hrs at 4°C. Wells were washed in Stellaris® Wash Buffer A with for 5 min at room temperature. Coverslips were transferred to a humidifying chamber and incubated with hybridization buffer (0.5 nM mRNA probe, 0.5 nM eRNA probe) for 14 hrs at 37°C.

Day 2: Coverslips were washed three times in Stellaris® Wash Buffer A for 30 min at 37°C. After a 5 min wash in Stellaris® Wash Buffer B at room temperature, coverslips were mounted using ProLong™ antifade with DAPI for imaging.

#### Quantification of Expression

A number of freely available programs have been developed to quantify smRNA FISH results. We used StarSearch (http://rajlab.seas.upenn.edu/StarSearch/launch.html), which was developed by Marshall J. Levesque and Arjun Raj at the University of Pennsylvania to automatically count individual RNAs. mRNA and eRNA detection involved two major steps. First, images for each probe set as well as a DAPI image are merged and cells were outlined. Punctae detection was carried out and additional adjustment of thresholds was performed. The same threshold range was used for all images, and this analysis was performed blind to treatment group. As a negative control, we quantified processed samples without FISH probes to determine non-specific background signals. Background signal for the Quasar® 570 channel (which was used to image *Fos* eRNA1 and *Gapdh* mRNA) was close to zero (0.3761 ± 0.07906 spots/cell). We did not detect any background spots in the Quasar® 670 channel, which was used to image *Fos* mRNA and *Gapdh* mRNA.

### RNA electrophoretic mobility shift assay (REMSA)

Mobility shift assays were conducted with synthetic Fluoresce- in-labeled ∼150-base RNA oligonucleotides and specified concentrations of recombinant CBP HAT domain (Sigma, 1319-1710) and CBP bromodomain (Abcam, ab198130). Oligonucleotides were synthesized using acRNA-containing Display plasmids as template DNA and the HiScribe T7 High Yield RNA synthesis kit (NEB) with Fluorescein-12-UTP (Sigma). RNA oligonucleotides (50nM) were heated to 95°C for 5min and refolded at room temperature prior to incubation with CBP protein (0-0.6M) in REMSA buffer (20mM HEPES, 40mM KCl, 1mM EDTA, 0.2mM DTT, 0.1mg ml-1 BSA, 0.1% Tween-20, and 20% Glycerol, and 0.1mg/ml tRNA(Sigma)) for 1hr at 37 °C. REMSA was performed using 1% agarose gel electrophoresis in 0.5xTBE. Electrophoretic mobility of Fluorescein-labeled RNA was assayed using fluorescence imaging on the Azure c600 Imaging System (Azure biosystems). RNA-CBP complex formation was quantified as the Fluorescein signal intensity appearing at the higher band (corresponding to CBP-bound RNA with lower electrophoretic mobility) divided by the total signal intensity (bound RNA plus free probe).

### Statistical Analysis

Required sample sizes were calculated using a freely available calculator (Lenth, R. V. (2006-9). Java Applets for Power and Sample Size [Computer software]. Available at http://www.stat.uiowa.edu/~rlenth/Power). Transcriptional differences from PCR experiments were compared with one-way ANOVA with post hoc tests where appropriate, two-way ANOVA with post hoc tests where appropriate, or Student’s t-tests/Mann-Whitney tests. Significance of smFISH data was assessed with Mann-Whitney test or Pearson correlation test. Statistical significance was designated at α = 0.05 for all analyses. Statistical and graphical analyses were performed with Graphpad software (Prism). Statistical assumptions (e.g., normality and homogeneity for parametric tests) were formally tested and boxplots were examined.

### Resource Availability

Sequencing data that support the findings of this study are available in Gene Expression Omnibus (GSE150499 and GSE150589). All relevant data that support the findings of this study are available on day-lab. org/resources or by request from the corresponding author (J.J.D.). CRISPR-Display constructs will be made available in the Addgene plasmid repository. R code TAPE analysis will be made available via GitLab (https://gitlab.rc.uab.edu/day-lab).

## Supplementary Data Tables

**Supplementary DataTable 1**. List and locations of all identified TAPEs

**Supplementary Data Table 2**. List and locations of all identified region-selective TAPEs

**Supplementary Data Table 3**. List a of all transcription factor binding motifs identified by HOMER analysis

**Supplementary Data Table 4**. Sequences of primers, guides, ASOs, sgRNAs, acRNAs, and smFISH probe sets

## Supplementary Figures

**Supplementary Figure 1.**
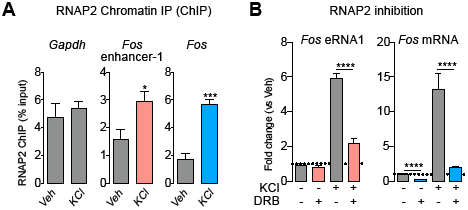
Activity dependence and synthesis of *Fos* eRNAs. **A**, RNAP2 ChIP reveals increased recruitment of RNAP2 to the *Fos* enhancer-1 and *Fos* gene body after KCl-mediated depolarization (unpaired t-test, for *Gapdh* promoter region *t*(6) 0.528, *p* = 0.6164; *Fos* enhancer-1 *t*(6) = 2.651, *p* = 0.038, and *Fos* gene body *t*(6) = 7.812, *p* = 0.0004). **B**, 2 hr pre-treatment with RNAP2 dependent transcription inhibitor DRB prior to 1 hr KCl treatment blocked KCl mediated induction of *Fos* eRNA1 and mRNA (two-way ANOVA, for eRNA1 *F*(1,42) = 27.84 *p* < 0.0001, and mRNA *F*(1,42) = 53.42 *p* < 0.0001, with Tukey’s post hoc test for multiple comparison). Data expressed as mean ± s.e.m. Multiple comparisons, \**p*<0.05, \*\**p*<0.01, \*\*\**p*<0.001, \*\*\*\**p*<0.0001.

**Supplementary Figure 2.**
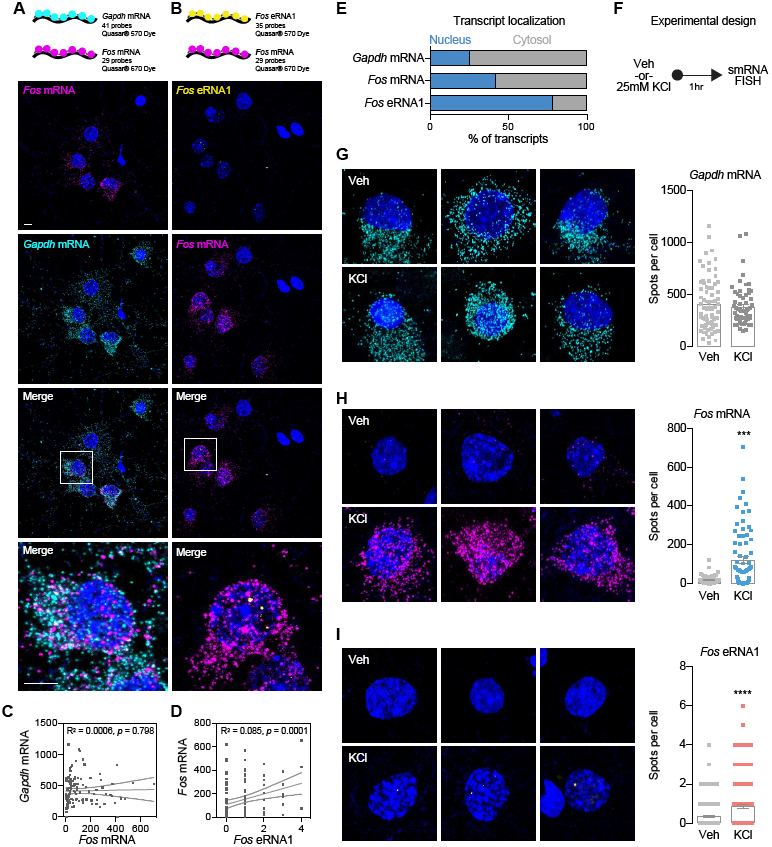
Cellular localization of *Fos* eRNA and mRNA. **A-B**, Top panel, illustration of smFISH probe sets indicating number of probes, dye, and LUT. Bottom panel, representative smFISH images for *Gapdh* mRNA (Quasar® 570) and *Fos* mRNA (Quasar® 670) (**A**), and *Fos* eRNA1 (Quasar® 570) and *Fos* mRNA (Quasar® 670) transcripts (**B**). Cell nuclei are stained with DAPI (blue), RNA transcripts are marked by smFISH probes (cyan, magenta, and yellow). Scale bar = 5 μm. **C-D**, Comparison and correlation of detected *Gapdh* mRNA, *Fos* mRNA, and *Fos* eRNA1 spots per cell. While there is no significant correlation between *Gapdh* mRNA and *Fos* mRNA (Pearson correlation for *Gapdh* mRNA and *Fos* mRNA, R2=0.000586, p=0.7982), *Fos* mRNA and *Fos* eRNA1 are positively correlated on a single cell level (Pearson correlation, R2=0.08481, p=0.0014). **E**, Compartmentalization of *Gapdh* mRNA, *Fos* mRNA, and *Fos* eRNA1. **F**, Experimental design for neuronal depolarization experiments. **G-I**, Representative images and summary data of *Gapdh* mRNA (top panel), *Fos* mRNA (middle panel), and eRNA1 (bottom panel) after 1 hr of Veh or 25mM KCl treatment. Number of detected *Fos* mRNA and *Fos* eRNA1 transcripts change significantly after stimulation (Mann-Whitney test for Gapdh n(veh)=72, n(KCl)=63, U=2187, p=0.7209; *Fos* mRNA n(veh)=77, n(KCl)=76, U=1929, p=0.0002; eRNA1 n(Veh)=124, n(KCl)=141, U=6540, p<0.0001). Data expressed as mean ± s.e.m. Multiple comparisons, *p<0.05, **p<0.01, ***p<0.001, ****p<0.0001.

**Supplementary Figure 3.**
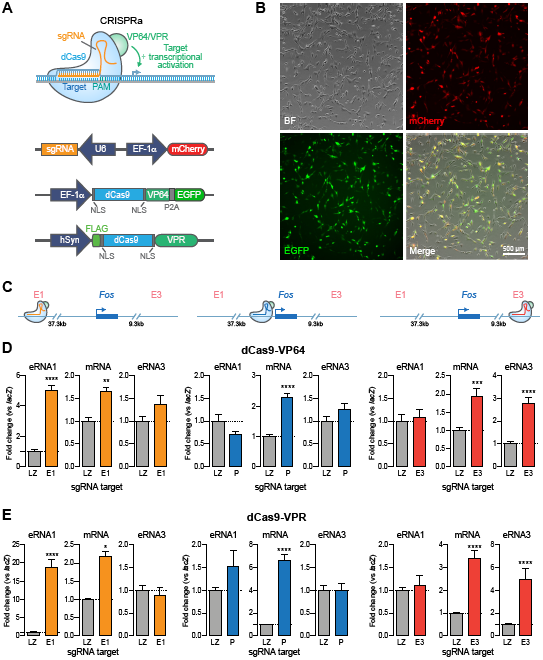
Enhancer activation increases *Fos* eRNA and mRNA expression. **A**, Illustration of CRISPR activation (CRISPRa) strategy for site-specific targeting of the transcriptional activator VP64 or VPR. **B**, C6 cells 16 hrs post nucleofection with VP64 containing plasmids (dCas-VP64 expression marked by GFP reporter, gRNA expression marked by mCherry reporter). **C**, sgRNA locations for *Fos* enhancer and promoter targeting. **D**, RT-qPCR analysis of VP64 mediated induction of *Fos* eRNAs and mRNA when targeted to individual sites surrounding the *Fos* gene, compared to the non-targeting *lacZ* control. CRISPRa resulted in site-specific upregulation of selected eRNAs and mRNA. Increasing *Fos* eRNA1 and eRNA3 levels resulted in increased *Fos* mRNA levels but not vice versa (*n* = 9 per group; one-way ANOVA for eRNA1 (*F*(4,40) = 66.22, *p* < 0.0001), eRNA3 (*F*(4,40) = 10.55, *p* < 0.0001), and mRNA (*F*(4,40) = 14.66, *p* < 0.0001); Dunnett’s multiple comparisons test). **E**, RT-qPCR analysis of VPR mediated induction of *Fos* eRNAs and mRNA when targeted to individual sites. CRISPRa resulted in site specific upregulation of selected eRNAs and mRNA compared to non-targeting *lacZ* control (*n* = 9 per group; one-way ANOVA for eRNA1 (*F*(4,40) = 49.47, *p* < 0.0001), eRNA3 (*F*(4,40) = 18.52, *p* < 0.0001), and mRNA (*F*(4,40) = 46.43, *p* < 0.0001); Dunnett’s multiple comparisons test. Increasing *Fos* eRNA1 and eRNA3 levels resulted in increased mRNA levels but not vice versa. Data expressed as mean ± s.e.m. Multiple comparisons, \**p* < 0.05, \*\**p* < 0.01, \*\*\**p* < 0.001, \*\*\*\**p* < 0.0001.

**Supplementary Figure 4.**
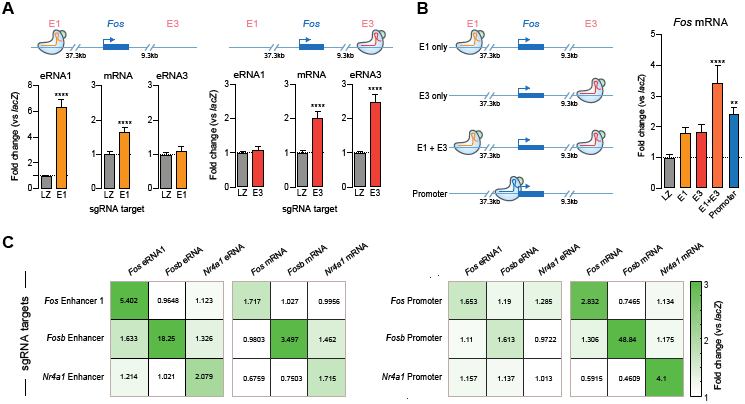
CRISPRa selectively activates targeted enhancer and linked gene without altering other enhancers or genes. **A**, CRISPRa targeting at a distal upstream enhancer (left) or a downstream enhancer (right) at the *Fos* gene locus. VPR targeting to enhancers induced robust eRNA transcription and also increased mRNA levels. Notably, *Fos* E1 targeting did not induce *Fos* eRNA3, and vice versa. Gene expression differences were measured with RT-qPCR (*n* = 18 per group; two-tailed Mann-Whitney test for all comparisons, *Fos* E1-eRNA3 *U* = 161, *p* = 0.9875; *Fos* P-eRNA1 *U* = 143, *p* = 0.5628; *Fos* P-eRNA3 *U* = 116, *p* = 0.1516; some data repeated from Fig.4). **B**, Multiplexed VPR-mediated enhancer activation in primary cortical neurons resulted in additive increases in *Fos* mRNA (*n* = 9 per group, Kurskal-Wallis *F*(4,40) = 25.04, *p* < 0.0001; Dunn’s multiple comparisons test). **C**, RT-qPCR data heatmap of CRISPRa experiments demonstrating specificity of enhancer (left) and promoter (right) activation. Enhancer activation induced eRNAs and mRNAs at the target genes with little effect on other tested eRNA or mRNAs. Promoter activation produced increases in mRNA with little effect on eRNA levels. Data expressed as mean ± s.e.m. Multiple comparisons, \**p* < 0.05, \*\**p* < 0.01, \*\*\**p* < 0.001, \*\*\*\**p*

## REFERENCES

Arner, E., Daub, C.O., Vitting-Seerup, K., Andersson, R., Lilje, B., Drablos, F., Lennartsson, A., Ronnerblad, M., Hrydziuszko, O., Vitezic, M., et al. (2015). Transcribed enhancers lead waves of coordinated transcription in transitioning mammalian cells. Science 347, 1010–1014.

Azofeifa, J. G., Allen, M. A., Hendrix, J. R., Read, T., Rubin, J. D., & Dowell, R. D. (2018). Enhancer RNA profiling predicts transcription factor activity. Genome Research, 28(3), 334–344. http://doi.org/10.1101/gr.225755.117

Baizabal, J.-M., Mistry, M., García, M. T., Gómez, N., Olukoya, O., Tran, D., et al. (2018). The Epigenetic State of PRDM16-Regulated Enhancers in Radial Glia Controls Cortical Neuron Position. Neuron, 99(1), 239–241. http://doi.org/10.1016/j.neuron.2018.06.031

Bose, D.A., Donahue, G., Reinberg, D., Shiekhattar, R., Bonasio, R., and Berger, S.L. (2017). RNA Binding to CBP Stimulates Histone Acetylation and Transcription. Cell 168, 135–149 e122.

Buenrostro, J. D., Wu, B., Chang, H. Y., & Greenleaf, W. J. (2015). ATAC-seq: A Method for Assaying Chromatin Accessibility Ge-nome-Wide. Current Protocols in Molecular Biology, 109(1), 21.29.1–21.29.9. http://doi.org/10.1002/0471142727.mb2129s109

Chavez, A., Scheiman, J., Vora, S., Pruitt, B.W., Tuttle, M., E, P.R.I., Lin, S., Kiani, S., Guzman, C.D., Wiegand, D.J., et al. (2015). Highly efficient Cas9-mediated transcriptional programming. Nat Methods 12, 326–328.

Chen, L.-F., Lin, Y. T., Gallegos, D. A., Hazlett, M. F., Gómez-Schiavon, M., Yang, M. G., et al. (2019). Enhancer Histone Acetylation Modulates Transcriptional Bursting Dynamics of Neuronal Activity-Inducible Genes. Cell Reports, 26(5), 1174–1188.e5. http://doi.org/10.1016/j.celrep.2019.01.032

Chen, X., Lepier, A., Berninger, B., Tolkovsky, A. M., & Herbert, J. (2012). Cultured subventricular zone progenitor cells transduced with neurogenin-2 become mature glutamatergic neurons and integrate into the dentate gyrus. PloS One, 7(2), e31547. http://doi.org/10.1371/journal.pone.0031547

Corces, M. R., Trevino, A. E., Hamilton, E. G., Greenside, P. G., Sinnott-Armstrong, N. A., Vesuna, S., et al. (2017). An improved ATAC-seq protocol reduces background and enables interrogation of frozen tissues. Nature Methods, 14(10), 959–962. http://doi.org/10.1038/nmeth.4396

Corradin, O., and Scacheri, P.C. (2014). Enhancer variants: evaluating functions in common disease. Genome Med 6, 85.

Davidson, S., Lear, M., Shanley, L., Hing, B., Baizan-Edge, A., Herwig, A., Quinn, J.P., Breen, G., McGuffin, P., Starkey, A., et al. (2011). Differential activity by polymorphic variants of a remote enhancer that supports galanin expression in the hypothalamus and amygdala: implications for obesity, depression and alcoholism. Neuropsychopharmacology 36, 2211–2221.

Day, J.J., Childs, D., Guzman-Karlsson, M.C., Kibe, M., Moulden, J., Song, E., Tahir, A., and Sweatt, J.D. (2013). DNA methylation regulates associative reward learning. Nat Neurosci 16, 1445–1452.

Di Ruscio, A., Ebralidze, A.K., Benoukraf, T., Amabile, G., Goff, L.A., Terragni, J., Figueroa, M.E., De Figueiredo Pontes, L.L., Alberich-Jorda, M., Zhang, P., et al. (2013). DNMT1-interacting RNAs block gene-specific DNA methylation. Nature 503, 371–376.

Djebali, S., Davis, C.A., Merkel, A., Dobin, A., Lassmann, T., Mortazavi, A., Tanzer, A., Lagarde, J., Lin, W., Schlesinger, F., et al. (2012). Landscape of transcription in human cells. Nature 489, 101–108.

Dobin, A., Davis, C. A., Schlesinger, F., Drenkow, J., Zaleski, C., Jha, S., et al. (2013). STAR: ultrafast universal RNA-seq aligner. Bioinformatics (Oxford, England), 29(1), 15–21. http://doi.org/ https://doi.org/10.1093/bioinformatics/bts635

Dong, X., Liao, Z., Gritsch, D., Hadzhiev, Y., Bai, Y., Locascio, J. J., et al. (2018). Enhancers active in dopamine neurons are a primary link between genetic variation and neuropsychiatric disease. Nature Neuroscience, 21(10), 1482–1492. http://doi.org/10.1038/s41593-018-0223-0

Eckart, N., Song, Q., Yang, R., Wang, R., Zhu, H., McCallion, A.S., and Avramopoulos, D. (2016). Functional Characterization of Schizophrenia-Associated Variation in CACNA1C. PLoS One 11, e0157086.

Edwards, A.C., Aliev, F., Bierut, L.J., Bucholz, K.K., Edenberg, H., Hesselbrock, V., Kramer, J., Kuperman, S., Nurnberger, J.I., Jr., Schuckit, M.A., et al. (2012). Genome-wide association study of comorbid depressive syndrome and alcohol dependence. Psychiatr Genet 22, 31–41.

Ehrman, L. A., Mu, X., Waclaw, R. R., Yoshida, Y., Vorhees, C. V., Klein, W. H., & Campbell, K. (2013). The LIM homeobox gene Isl1 is required for the correct development of the striatonigral pathway in the mouse. Proceedings of the National Academy of Sciences of the United States of America, 110(42), E4026–35. http://doi.org/10.1073/pnas.1308275110

Flavell, S. W., Cowan, C. W., Kim, T.-K., Greer, P. L., Lin, Y., Paradis, S., et al. (2006). Activity-Dependent Regulation of MEF2 Transcription Factors Suppresses Excitatory Synapse Number. Science (New York, N.Y.), 311(5763), 1008–1012. http://doi.org/10.1126/science.1122511

Fleischmann, A., Hvalby, O., Jensen, V., Strekalova, T., Zacher, C., Layer, L.E., Kvello, A., Reschke, M., Spanagel, R., Sprengel, R., et al. (2003). Impaired long-term memory and NR2A-type NMDA receptor-dependent synaptic plasticity in mice lacking c-Fos in the CNS. J Neurosci 23, 9116–9122.

Frank, C.L., Liu, F., Wijayatunge, R., Song, L., Biegler, M.T., Yang, M.G., Vockley, C.M., Safi, A., Gersbach, C.A., Crawford, G.E., et al. (2015). Regulation of chromatin accessibility and Zic binding at enhancers in the developing cerebellum. Nat Neurosci 18, 647–656.

Fullard, J. F., Hauberg, M. E., Bendl, J., Egervari, G., Cirnaru, M.-D., Reach, S. M., et al. (2018). An atlas of chromatin accessibility in the adult human brain. Genome Research, 28(8), 1243–1252. http://doi.org/10.1101/gr.232488.117

Furlong, E. E. M., & Levine, M. (2018). Developmental enhancers and chromosome topology. Science (New York, N.Y.), 361(6409), 1341–1345. http://doi.org/10.1126/science.aau0320

Galichet, C., Guillemot, F., & Parras, C. M. (2008). Neurogenin 2 has an essential role in development of the dentate gyrus. Development (Cambridge, England), 135(11), 2031–2041. http://doi.org/10.1242/dev.015115

Gallegos, D. A., Chan, U., Chen, L.-F., & West, A. E. (2018). Chromatin Regulation of Neuronal Maturation and Plasticity. Trends in Neurosciences, 41(5), 311–324. http://doi.org/10.1016/j.tins.2018.02.009

Gordon, C.T., and Lyonnet, S. (2014). Enhancer mutations and phenotype modularity. Nat Genet 46, 3–4.

Gray, J.M., Kim, T.K., West, A.E., Nord, A.S., Markenscoff-Papadimitriou, E., and Lomvardas, S. (2015). Genomic Views of Transcriptional Enhancers: Essential Determinants of Cellular Identity and Activity-Dependent Responses in the CNS. J Neurosci 35, 13819–13826.

Hamilton, W. B., Mosesson, Y., Monteiro, R. S., Emdal, K. B., Knudsen, T. E., Francavilla, C., et al. (2019). Dynamic lineage priming is driven via direct enhancer regulation by ERK. Nature, 575(7782), 355–360. http://doi.org/10.1038/s41586-019-1732-z

Hangauer, M.J., Vaughn, I.W., and McManus, M.T. (2013). Pervasive transcription of the human genome produces thousands of previously unidentified long intergenic noncoding RNAs. PLoS Genet 9, e1003569.

Heinz, S., Romanoski, C.E., Benner, C., and Glass, C.K. (2015). The selection and function of cell type-specific enhancers. Nat Rev Mol Cell Biol 16, 144–154.

Hendrickson, D., Kelley, D.R., Tenen, D., Bernstein, B., and Rinn, J.L. (2016). Widespread RNA binding by chromatin-associated proteins. Genome Biol 17, 28.

Hilton, I.B., D’Ippolito, A.M., Vockley, C.M., Thakore, P.I., Crawford, G.E., Reddy, T.E., and Gersbach, C.A. (2015). Epigenome editing by a CRISPR-Cas9-based acetyltransferase activates genes from promoters and enhancers. Nat Biotechnol 33, 510–517.

Hnisz, D., Abraham, B.J., Lee, T.I., Lau, A., Saint-Andre, V., Sigova, A.A., Hoke, H.A., and Young, R.A. (2013). Super-enhancers in the control of cell identity and disease. Cell 155, 934–947.

Hsieh, C.L., Fei, T., Chen, Y., Li, T., Gao, Y., Wang, X., Sun, T., Sweeney, C.J., Lee, G.S., Chen, S., et al. (2014). Enhancer RNAs participate in androgen receptor-driven looping that selectively enhances gene activation. Proc Natl Acad Sci U S A 111, 7319–7324.

Impey, S., McCorkle, S. R., Cha-Molstad, H., Dwyer, J. M., Yochum, G. S., Boss, J. M., et al. (2004). Defining the CREB Regulon: A Genome-Wide Analysis of Transcription Factor Regulatory Regions. Cell, 119(7), 1041–1054. http://doi.org/10.1016/j.cell.2004.10.032

Inoue, Y.U., and Inoue, T. (2016). Brain enhancer activities at the gene-poor 5p14.1 autism-associated locus. Sci Rep 6, 31227.

Jeong, Y., Leskow, F.C., El-Jaick, K., Roessler, E., Muenke, M., Yocum, A., Dubourg, C., Li, X., Geng, X., Oliver, G., et al. (2008). Regulation of a remote Shh forebrain enhancer by the Six3 homeoprotein. Nat Genet 40, 1348–1353.

Joo, J.Y., Schaukowitch, K., Farbiak, L., Kilaru, G., and Kim, T.K. (2016). Stimulus-specific combinatorial functionality of neuronal c-fos enhancers. Nat Neurosci 19, 75–83.

Kaikkonen, M.U., Spann, N.J., Heinz, S., Romanoski, C.E., Allison, K.A., Stender, J.D., Chun, H.B., Tough, D.F., Prinjha, R.K., Benner, C., et al. (2013). Remodeling of the enhancer landscape during macrophage activation is coupled to enhancer transcription. Mol Cell 51, 310–325.

Kearns, N.A., Pham, H., Tabak, B., Genga, R.M., Silverstein, N.J., Garber, M., and Maehr, R. (2015). Functional annotation of native enhancers with a Cas9-histone demethylase fusion. Nat Methods 12, 401–403.

Kim, T.K., Hemberg, M., and Gray, J.M. (2015). Enhancer RNAs: a class of long noncoding RNAs synthesized at enhancers. Cold Spring Harb Perspect Biol 7, a018622.

Kim, T.K., Hemberg, M., Gray, J.M., Costa, A.M., Bear, D.M., Wu, J., Harmin, D.A., Laptewicz, M., Barbara-Haley, K., Kuersten, S., et al. (2010). Widespread transcription at neuronal activity-regulated enhancers. Nature 465, 182–187.

Kim, T.K., and Shiekhattar, R. (2015). Architectural and Functional Commonalities between Enhancers and Promoters. Cell 162, 948–959.

Konermann, S., Brigham, M.D., Trevino, A.E., Joung, J., Abudayyeh, O.O., Barcena, C., Hsu, P.D., Habib, N., Gootenberg, J.S., Nishimasu, H., et al. (2015). Genome-scale transcriptional activation by an engineered CRISPR-Cas9 complex. Nature 517, 583–588.

Köster, J., & Rahmann, S. (2018). Snakemake-a scalable bioinformatics workflow engine. Bioinformatics (Oxford, England), 34(20), 3600–3600. http://doi.org/10.1093/bioinformatics/bty350

Kwan, K. Y., Lam, M. M. S., Krsnik, Z., Kawasawa, Y. I., Lefebvre, V., & Sestan, N. (2008). SOX5 postmitotically regulates migration, postmigratory differentiation, and projections of subplate and deep-layer neocortical neurons. Proceedings of the National Academy of Sciences of the United States of America, 105(41), 16021–16026. http://doi.org/10.1073/pnas.0806791105

Lai, F., Orom, U.A., Cesaroni, M., Beringer, M., Taatjes, D.J., Blobel, G.A., and Shiekhattar, R. (2013). Activating RNAs associate with Mediator to enhance chromatin architecture and transcription. Nature 494, 497–501.

Langmead, B., & Salzberg, S. L. (2012). Fast gapped-read alignment with Bowtie 2. Nature Methods, 9(4), 357–359. http://doi.org/10.1038/nmeth.1923

Leighton, P.A., Saam, J.R., Ingram, R.S., Stewart, C.L., and Tilghman, S.M. (1995). An enhancer deletion affects both H19 and Igf2 expression. Genes & development 9, 2079–2089.

Li, H., Handsaker, B., Wysoker, A., Fennell, T., Ruan, J., Homer, N., et al. (2009). The Sequence Alignment/Map format and SAMtools. Bioinformatics (Oxford, England), 25(16), 2078–2079. http://doi.org/10.1093/bioinformatics/btp352

Li, K., Liu, Y., Cao, H., Zhang, Y., Gu, Z., Liu, X., et al. (2020). Interrogation of enhancer function by enhancer-targeting CRISPR epigenetic editing. Nature Communications, 11(1), 485–16. http://doi.org/10.1038/s41467-020-14362-5

Li, P., Marshall, L., Oh, G., Jakubowski, J. L., Groot, D., He, Y., et al. (2019b). Epigenetic dysregulation of enhancers in neurons is associated with Alzheimer’s disease pathology and cognitive symptoms. Nature Communications, 10(1), 15056–14. http://doi.org/10.1038/s41467-019-10101-7

Li, W., Notani, D., Ma, Q., Tanasa, B., Nunez, E., Chen, A.Y., Merkurjev, D., Zhang, J., Ohgi, K., Song, X., et al. (2013). Functional roles of enhancer RNAs for oestrogen-dependent transcriptional activation. Nature 498, 516–520.

Li, W., Notani, D., and Rosenfeld, M.G. (2016). Enhancers as non-coding RNA transcription units: recent insights and future perspectives. Nat Rev Genet 17, 207–223.

Liu, S.J., Horlbeck, M.A., Cho, S.W., Birk, H.S., Malatesta, M., He, D., Attenello, F.J., Villalta, J.E., Cho, M.Y., Chen, Y., et al. (2017). CRISPRi-based genome-scale identification of functional long noncoding RNA loci in human cells. Science 355.

Liu, S.J., Nowakowski, T.J., Pollen, A.A., Lui, J.H., Horlbeck, M.A., Attenello, F.J., He, D., Weissman, J.S., Kriegstein, A.R., Diaz, A.A., et al. (2016). Single-cell analysis of long non-coding RNAs in the developing human neocortex. Genome Biol 17, 67.

Lopes, R., Korkmaz, G., and Agami, R. (2016). Applying CRIS-PR-Cas9 tools to identify and characterize transcriptional enhancers. Nat Rev Mol Cell Biol 17, 597–604.

Malik, A.N., Vierbuchen, T., Hemberg, M., Rubin, A.A., Ling, E., Couch, C.H., Stroud, H., Spiegel, I., Farh, K.K., Harmin, D.A., et al. (2014). Genome-wide identification and characterization of functional neuronal activity-dependent enhancers. Nat Neurosci 17, 1330–1339.

Mikhaylichenko, O., Bondarenko, V., Harnett, D., Schor, I. E., Males, M., Viales, R. R., & Furlong, E. E. M. (2018). The degree of enhancer or promoter activity is reflected by the levels and directionality of eRNA transcription. Genes & Development, 32(1), 42–57. http://doi.org/10.1101/gad.308619.117

Moffat, J., Grueneberg, D. A., Yang, X., Kim, S. Y., Kloepfer, A. M., Hinkle, G., et al. (2006). A lentiviral RNAi library for human and mouse genes applied to an arrayed viral high-content screen. Cell, 124(6), 1283–1298. http://doi.org/10.1016/j.cell.2006.01.040

Network, and Pathway Analysis Subgroup of Psychiatric Genomics, C. (2015). Psychiatric genome-wide association study analyses implicate neuronal, immune and histone pathways. Nat Neurosci 18, 199–209.

Nord, A.S., Blow, M.J., Attanasio, C., Akiyama, J.A., Holt, A., Hosseini, R., Phouanenavong, S., Plajzer-Frick, I., Shoukry, M., Afzal, V., et al. (2013). Rapid and pervasive changes in genome-wide enhancer usage during mammalian development. Cell 155, 1521–1531.

Nott, A., Holtman, I. R., Coufal, N. G., Schlachetzki, J. C. M., Yu, M., Hu, R., et al. (2019). Brain cell type-specific enhancer-promoter interactome maps and disease-risk association. Science (New York, N.Y.), 366(6469), 1134–1139. http://doi.org/10.1126/science.aay0793

Pattabiraman, K., Golonzhka, O., Lindtner, S., Nord, A.S., Taher, L., Hoch, R., Silberberg, S.N., Zhang, D., Chen, B., Zeng, H., et al. (2014). Transcriptional regulation of enhancers active in protodomains of the developing cerebral cortex. Neuron 82, 989–1003.

Quinlan, A. R., & Hall, I. M. (2010). BEDTools: a flexible suite of utilities for comparing genomic features. Bioinformatics (Oxford, England), 26(6), 841–842. http://doi.org/10.1093/bioinformatics/btq033

Quinn, J.J., and Chang, H.Y. (2016). Unique features of long non-coding RNA biogenesis and function. Nat Rev Genet 17, 47–62.

Rahnamoun, H., Lee, J., Sun, Z., Lu, H., Ramsey, K. M., Komives, E. A., & Lauberth, S. M. (2018). RNAs interact with BRD4 to promote enhanced chromatin engagement and transcription activation. Nature Structural & Molecular Biology, 25(8), 687–697. http://doi.org/10.1038/s41594-018-0102-0

Rinn, J.L., and Chang, H.Y. (2012). Genome regulation by long noncoding RNAs. Annu Rev Biochem 81, 145–166.

Robson, M. I., Ringel, A. R., & Mundlos, S. (2019). Regulatory Landscaping: How Enhancer-Promoter Communication Is Sculpted in 3D. Molecular Cell, 74(6), 1110–1122. http://doi.org/10.1016/j.molcel.2019.05.032

Roussos, P., Mitchell, A.C., Voloudakis, G., Fullard, J.F., Pothula, V.M., Tsang, J., Stahl, E.A., Georgakopoulos, A., Ruderfer, D.M., Charney, A., et al. (2014). A role for noncoding variation in schizophrenia. Cell Rep 9, 1417–1429.

Sanjana, N.E., Shalem, O., and Zhang, F. (2014). Improved vectors and genome-wide libraries for CRISPR screening. Nat Methods 11, 783–784.

Sanjana, N.E., Wright, J., Zheng, K., Shalem, O., Fontanillas, P., Joung, J., Cheng, C., Regev, A., and Zhang, F. (2016). High-resolution interrogation of functional elements in the noncoding genome. Science 353, 1545–1549.

Sanyal, A., Lajoie, B.R., Jain, G., and Dekker, J. (2012). The longrange interaction landscape of gene promoters. Nature 489, 109–113.

Sanchez-Mut, J. V., Heyn, H., Silva, B. A., Dixsaut, L., Garcia-Esparcia, P., Vidal, E., et al. (2018). PM20D1 is a quantitative trait locus associated with Alzheimer’s disease, 24(5), 598–603. http://doi.org/10.1038/s41591-018-0013-y

Savell, K. E., Bach, S. V., Zipperly, M. E., Revanna, J. S., Goska, N. A., Tuscher, J. J., et al. (2018). A neuron-optimized CRISPR/dCas9 activation system for robust and specific gene regulation. bioRxiv, 371500. http://doi.org/10.1101/371500

Savell, K.E., Gallus, N.V., Simon, R.C., Brown, J.A., Revanna, J.S., Osborn, M.K., Song, E.Y., O’Malley, J.J., Stackhouse, C.T., Norvil, A., et al. (2016). Extra-coding RNAs regulate neuronal DNA methylation dynamics. Nat Commun 7, 12091.

Savell, K., Sultan, F., & Day, J. (2019). A Novel Dual Lentiviral CRISPR-based Transcriptional Activation System for Gene Expression Regulation in Neurons. Bio-Protocol, 9(17). http://doi.org/10.21769/BioProtoc.3348

Scharer, C. D., Blalock, E. L., Barwick, B. G., Haines, R. R., Wei, C., Sanz, I., & Boss, J. M. (2016). ATAC-seq on biobanked specimens defines a unique chromatin accessibility structure in naïve SLE B cells. Scientific Reports, 6(1), 27030–9. http://doi.org/10.1038/srep27030

Schaukowitch, K., Joo, J.Y., Liu, X., Watts, J.K., Martinez, C., and Kim, T.K. (2014). Enhancer RNA facilitates NELF release from immediate early genes. Mol Cell 56, 29–42.

Schizophrenia Working Group of the Psychiatric Genomics, C. (2014). Biological insights from 108 schizophrenia-associated genetic loci. Nature 511, 421–427.

Shechner, D.M., Hacisuleyman, E., Younger, S.T., and Rinn, J.L. (2015). Multiplexable, locus-specific targeting of long RNAs with CRISPR-Display. Nat Methods 12, 664–670.

Sigova, A.A., Abraham, B.J., Ji, X., Molinie, B., Hannett, N.M., Guo, Y.E., Jangi, M., Giallourakis, C.C., Sharp, P.A., and Young, R.A. (2015). Transcription factor trapping by RNA in gene regulatory elements. Science 350, 978–981.

Spieler, D., Kaffe, M., Knauf, F., Bessa, J., Tena, J.J., Giesert, F., Schormair, B., Tilch, E., Lee, H., Horsch, M., et al. (2014). Restless legs syndrome-associated intronic common variant in Meis1 alters enhancer function in the developing telencephalon. Genome research 24, 592–603.

Song, M., Yang, X., Ren, X., Maliskova, L., Li, B., Jones, I. R., et al. (2019). Mapping cis-regulatory chromatin contacts in neural cells links neuropsychiatric disorder risk variants to target genes. Nature Genetics, 51(8), 1252–1262. http://doi.org/10.1038/s41588-019-0472-1

Telese, F., Ma, Q., Perez, P.M., Notani, D., Oh, S., Li, W., Comoletti, D., Ohgi, K.A., Taylor, H., and Rosenfeld, M.G. (2015). LRP8-Reelin-Regulated Neuronal Enhancer Signature Underlying Learning and Memory Formation. Neuron 86, 696–710.

Tyssowski, K. M., DeStefino, N. R., Cho, J.-H., Dunn, C. J., Poston, R. G., Carty, C. E., et al. (2018). Different Neuronal Activity Patterns Induce Different Gene Expression Programs. Neuron, 98(3), 530–546.e11. http://doi.org/10.1016/j.neuron.2018.04.001

Vermunt, M.W., Reinink, P., Korving, J., de Bruijn, E., Creyghton, P.M., Basak, O., Geeven, G., Toonen, P.W., Lansu, N., Meunier, C., et al. (2014). Large-scale identification of coregulated enhancer networks in the adult human brain. Cell Rep 9, 767–779.

Voisin, S., Almen, M.S., Zheleznyakova, G.Y., Lundberg, L., Zarei, S., Castillo, S., Eriksson, F.E., Nilsson, E.K., Bluher, M., Bottcher, Y., et al. (2015). Many obesity-associated SNPs strongly associate with DNA methylation changes at proximal promoters and enhancers. Genome Med 7, 103.

Wang, D., Garcia-Bassets, I., Benner, C., Li, W., Su, X., Zhou, Y., Qiu, J., Liu, W., Kaikkonen, M.U., Ohgi, K.A., et al. (2011). Reprogramming transcription by distinct classes of enhancers functionally defined by eRNA. Nature 474, 390–394.

Wang, Z., Chivu, A. G., Choate, L. A., Rice, E. J., Miller, D. C., Chu, T., et al. (2020). Accurate imputation of histone modifications using transcription. bioRxiv, 2, 2020.04.08.032730. http://doi.org/10.1101/2020.04.08.032730

Yao, P., Lin, P., Gokoolparsadh, A., Assareh, A., Thang, M.W., and Voineagu, I. (2015). Coexpression networks identify brain region-specific enhancer RNAs in the human brain. Nat Neurosci 18, 1168–1174.

Zhang, Y., Liu, T., Meyer, C. A., Eeckhoute, J., Johnson, D. S., Bernstein, B. E., et al. (2008). Model-based analysis of ChIP-Seq (MACS). Genome Biology, 9(9), R137–9. http://doi.org/10.1186/gb-2008-9-9-r137

Zipperly, M. E., Sultan, F. A., Graham, G.-E., Brane, A. C., Simpkins, N. A., Ianov, L., & Day, J. J. (2020). Regulation of dopamine-dependent transcription and cocaine action by Gadd45b. bioRxiv, 2020.05.01.072926. http://doi.org/10.1101/2020.05.01.072926

Zovkic, I.B., Paulukaitis, B.S., Day, J.J., Etikala, D.M., and Sweatt, J.D. (2014). Histone H2A.Z subunit exchange controls consolidation of recent and remote memory. Nature 515, 582–586.

